# A Hybrid Gag Lattice as a Structural Intermediate in HIV-1 Maturation

**DOI:** 10.64898/2026.01.10.698794

**Authors:** Megan E. Meuser, Chunxiang Wu, Amanda C. Macke, Jiong Shi, Jing Cheng, Rachel R. Yang, Christian Freniere, Jianfeng Lin, Christopher Aiken, Juan R. Perilla, Yong Xiong

**Affiliations:** Department of Molecular Biophysics and Biochemistry, Yale University, New Haven, CT, USA; Department of Chemistry and Biochemistry, University of Delaware, Newark, DE, USA; Department of Pathology, Microbiology and Immunology, Vanderbilt University Medical Center, Nashville, TN, USA

**Author notes:** Correspondence (Y.X.). Denotes equal contributions (Megan E. Meuser, Chunxiang Wu).

## Abstract

HIV-1 maturation converts the spherical immature Gag lattice into the conical mature capsid required for infectivity, yet the structural route linking these two lattice endpoints remains unclear. Using single-particle cryo-electron microscopy and cryo-electron tomography on reconstituted assemblies and virus-like particles (VLPs), we identify a distinct hybrid lattice in which the capsid protein (CA) N-terminal domain adopts a mature-like conformation, whereas the CA C-terminal domain–SP1 layer remains immature. This architecture is observed in native, enveloped VLPs produced in human cells, demonstrating formation under physiologically relevant conditions. The hybrid lattice engages three myo-inositol hexakisphosphate (IP6) molecules per CA–SP1 hexamer, underscoring IP6-dependent stabilization; accordingly, excess IP6 enriches the hybrid population. Tomographic mapping shows that hybrid regions coexist with the immature lattice within the same particle and are enriched near lattice discontinuities, consistent with edge-localized remodeling that can accommodate conformational rearrangements and partial disassembly and reassembly. Disrupting a hybrid-specific inter-hexamer contact preserves immature lattice assembly and particle release but abrogates infectivity, compromises core integrity, and prevents mature lattice formation, implicating the hybrid architecture as an on-pathway intermediate. Molecular dynamics simulations further support coordinated rearrangements that bias the system away from the immature configuration toward hybrid and then mature organizations. Together, these results support a maturation model in which localized displacive remodeling and partial disassembly/reassembly act in concert, with IP6 tuning the balance among lattice states. This study provides new insight into HIV-1 maturation and identifies the hybrid lattice as a potential therapeutic target.

## Introduction

HIV-1 maturation is an essential process in the viral life cycle that enables the production of infectious virions. During maturation, the viral protease cleaves the Gag polyprotein into its individual subunits: matrix (MA), capsid (CA), spacer peptide 1 (SP1), nucleocapsid (NC), spacer peptide 2 (SP2), and p6 domains. These specific and tightly regulated proteolytic cleavage events transform the immature Gag lattice into a mature capsid in the fully infectious virion^1^. Disrupting maturation yields largely noninfectious virions^2–4^, underscoring its central role and the need to define its mechanism. Defining this mechanism will fill a key gap in HIV-1 biology and reveal therapeutic targets to block production of infectious virions.

HIV-1 maturation is currently framed by two distinct states of structural lattices: (1) the immature lattice, which is formed by Gag polyproteins or early proteolytic products; and (2) the mature capsid, which is composed of cleaved CA proteins. CA, composed of an N-terminal domain (NTD) and C-terminal domain (CTD) connected by a flexible linker, serves as a key structural component in both the immature lattice and the mature capsid. In the immature lattice, the CA-SP1 domain of Gag organizes into a lattice of hexameric subunits in an incomplete spherical structure^5–7^. Residues at the CA-SP1 junction facilitate the formation of a six-helix bundle necessary for immature lattice stabilization^8–10^. Additionally, the immature lattice engages one IP6 molecule per CA–SP1 hexamer, coordinated by six pairs of K158 and K227 residues^11–16^. This interaction plays a critical role in stabilizing the immature lattice and enriching IP6 within HIV-1 virions, ensuring proper CA lattice maturation^12, 16^. Upon maturation, the mature capsid consists of approximately 250 hexameric and 12 pentameric CA capsomeres^17^, with each coordinating two IP6 molecules in the central pore between residues R18 and K25^15, 18^. Capsomeres are stabilized internally by CA_NTD_-CA_NTD_ and CA_NTD_-CA_CTD_ interactions, while CA_CTD_-CA_CTD_ interactions link adjacent capsomeres to form the cone-shaped mature capsid. The substantial differences in lattice organization and specific protein-protein interactions between the immature and mature forms necessitate a complete rearrangement of CA-CA interfaces to accommodate the structural transition.

Although the structures of the immature Gag lattice and mature capsid have been very thoroughly characterized, the transition between the two distinct structures remains elusive. Three models have been proposed for this transition process: (i) de novo reassembly, where Gag proteolytic cleavage releases CA into the viral lumen, allowing mature capsid formation from CA monomers; (ii) displacive reorganization, in which the immature lattice transitions directly into the mature CA lattice without disassembly; (iii) a combination of the two (**Fig. 1A**). Support for a disassembly/reassembly mechanism primarily comes from biochemical measurements showing that HIV-1 virions contain a substantial pool of soluble CA that is not incorporated into the mature core^19–21^, implying that CA molecules dissociate before reassembly. Conversely, several observations are consistent with displacive lattice remodeling without extensive CA release. Protease cleavage of *in vitro* assembled CA-NC tubes, which are already in a mature-like arrangement, can nonetheless drive large-scale coordinated rearrangements of CA_NTD_ and CA_CTD_ without major lattice disassembly, highlighting the intrinsic structural plasticity of the CA lattice^22^. In addition, cryo-ET studies have revealed multiple mature-sized cores within a single viral membrane, supporting models in which capsid formation proceeds through a rolling-sheet, displacive transformation rather than complete de novo reassembly^23,24^. Furthermore, *in vitro* protease processing of Gag virus-like particles (VLPs) have demonstrated that while a fraction of CA and CA-SP1 dissociates from the lattice, a substantial portion remains lattice-bound, suggesting that maturation involves a coordinated local displacive rearrangement and partial CA release rather than a purely sequential mechanism^25,26^. However, the dense packing of CA within the immature or mature lattice imposes substantial geometric and energetic constraints on conformational transitions, and a discrete structural intermediate bridging the states has yet to be observed. In a recent study of the capsid inhibitor Lenacapavir (LEN) bound to the immature Gag lattice, LEN binding induced an ∼8 degrees rotation of CA_NTD_^27^, revealing measurable conformational flexibility within the Gag lattice. This finding suggests that the CA domain is pliable and capable of local conformational remodeling between different structural states, a plasticity that may be important for capsid maturation.

**Figure 1.**
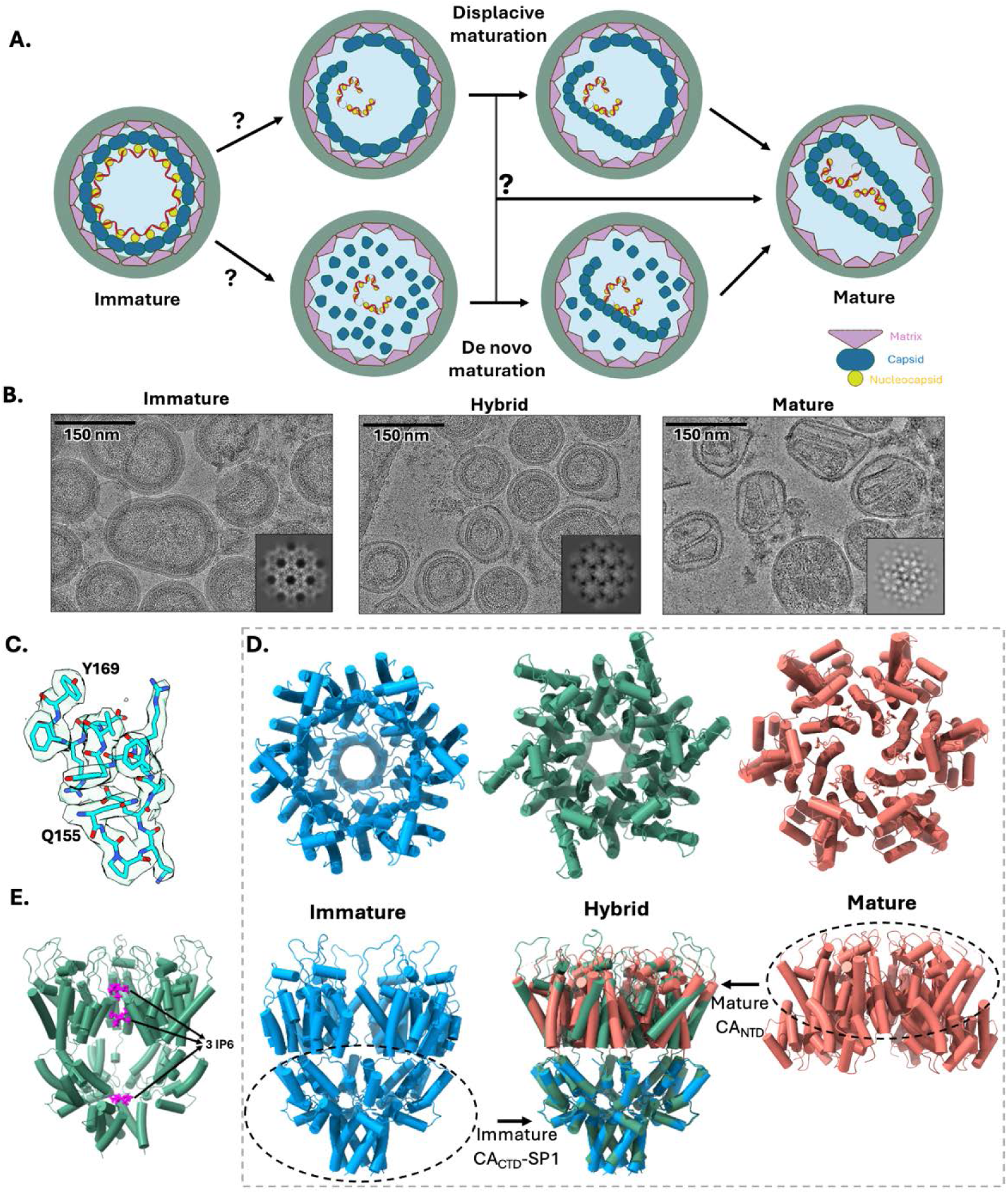
Identification of a hybrid HIV-1 Gag lattice assembly. A. Cartoon schematic representing the potential HIV-1 maturation pathways. B. Representative cryo-EM micrographs of immature (NL-MA/NC, left), hybrid (NL-CA/SP1, middle), and mature (R9 ΔEnv, right) VLPs. A corresponding 2D class average from each dataset is shown in the bottom-right corner. C. Example of cryo-EM density for selected residues from the hybrid lattice. D. Structural comparison of immature (blue), hybrid (green), and mature (coral) CA-SP1 and CA hexamers shown from the top view (top) and side view (bottom). The mature CA_NTD_ and immature CA_CTD_ are aligned to the hybrid hexamer (center bottom) to highlight structural similarities. E. Side view of a hybrid hexamer showing three bound IP6 molecules (magenta): two within the CA_NTD_ binding sites and one within the CA_CTD_-SP1 binding site.

To investigate the structural transitions during HIV-1 capsid maturation, we performed comprehensive single-particle cryo-electron microscopy (cryo-EM) analysis on *in vitro* lattice assemblies and *in situ* membrane-enveloped VLPs at different stages of maturation. Strikingly, our analysis revealed a hybrid lattice featuring both mature-like and immature-like features, providing the first structural evidence for a potential intermediate state of maturation. *In situ* cryo-EM analysis confirmed that this hybrid lattice type exists in VLPs produced from mammalian cells. We further found that exposure to elevated IP6 concentrations can enhance the formation of this hybrid lattice, corroborating the critical role of IP6 in modulating lattice organization. Cryo-electron tomography (cryo-ET) of VLPs revealed hybrid lattices enriched near lattice discontinuities, suggesting local structural transition zones. Furthermore, mutations designed to specifically destabilize the hybrid lattice impaired capsid function without preventing immature lattice formation, suggesting that the hybrid lattice is an on-pathway intermediate during the later stages of capsid remodeling. Together, our findings support a combined-pathway model of HIV-1 maturation: partial CA disassembly releases IP6, and the resulting excess IP6 promotes hybrid-lattice formation, seeding subsequent structural transitions.

## Results

### High-resolution structural analysis of the hybrid CA lattice assembly

HIV-1 maturation involves the transformation from a spherical immature Gag lattice to a conical mature form of the viral capsid (**Fig. 1A**). Despite extensive study, the intermediate steps linking these states remain poorly defined. To clarify the mechanism of maturation, we employed cryo-EM methods on both *in vitro* lattice assemblies and *in situ* VLPs at multiple stages of the maturation process (**Fig. 1B**). This analysis revealed a previously uncharacterized hybrid CA lattice in several types of specimens. Specifically, we obtained high-resolution cryo-EM maps of the hybrid lattice in *in vitro* assembled CA-NC (2.9 Å resolution), CA-SP1_T8I_ (2.6 Å resolution), and CA-SP1_BVM_ (3.7 Å resolution) constructs as well as *in situ* cryo-EM maps for VLPs including CA-NC (3.2 Å resolution) and CA-SP1 (3.9 Å resolution) (**Table S1**, **Table S2**, **Fig. 1C and Fig. S1**).

Our results revealed that the hybrid lattice structure exhibits a mature-like CA_NTD_ arrangement while retaining an immature CA_CTD_-SP1 configuration, potentially reflecting an intermediate stage in the maturation process (**Fig. 1D**). The hybrid lattice also incorporates three copies of IP6 molecules in each hexamer (**Fig. 1E**). This includes: (1) two IP6 molecules associated with the mature-like CA_NTD_ hexamers, coordinated by R18 and K25; and (2) one IP6 molecule within the immature-identical CA_CTD_-SP1 hexamers, coordinated by K158 and K227. These results indicate that IP6 may play a regulatory role in stabilizing the hybrid conformation. Furthermore, the hybrid lattice appears to capture a critical intermediate state between the lattice assembly patterns of the immature and mature lattices. We hypothesize that this structure represents a transition state within the displacive model of maturation, providing high-resolution structural evidence for this mechanism.

We initially observed the hybrid lattice using *in vitro* lattice assemblies from a CA-NC construct. In previous studies, CA-NC constructs were frequently designed with N-terminal extensions to prevent the formation of the CA N-terminal β-hairpin, a critical structure for the mature capsid^28^, and were often combined with stabilizing mutations to further favor the immature lattice conformation^8–10, 15, 29–32^. Instead, we employed the wild-type CA with a natural N-terminus and assembled the lattice in the presence of nucleic acids and IP6. Under these conditions, the previously unobserved hybrid lattice was revealed in our cryo-EM studies. This finding differs from many CA-NC-based reconstitution studies, which predominantly yield immature lattices, a bias that may arise from experimental design features, such as residual MA sequences or artificial N-terminal tags, that extend the CA N-terminus and can disfavor the formation of the mature β-hairpin^8–10, 15, 29–32^. Model building and thorough structural analysis of our newly observed lattice revealed a hybrid assembly pattern that integrates features of both the immature Gag and mature CA lattices. Specifically, in the hybrid assembly, CA_CTD_-SP1 adopts a conformation nearly identical to that of the immature Gag lattice (RMSD 0.4 Å), while CA_NTD_ adopts a fold resembling that in the mature CA lattice (RMSD 0.7 Å) (**Fig. 1D and Fig. S2**). The identification of this hybrid lattice provides direct structural evidence of a potential intermediate state, bridging the gap between immature and mature assemblies, and offers new insights into the possible mechanisms governing HIV-1 maturation.

### *In situ* cryo-EM and cryo-ET reveal hybrid lattice formation at lattice discontinuities and its promotion by IP6

Having identified the hybrid lattice *in vitro*, we next sought to determine whether it could be detected *in situ* under native conditions. We analyzed mammalian-produced CA-NC and CA-SP1 VLPs, which have a cleavage-resistant CA-NC or CA-SP1 construct of Gag. These VLPs were intact and vitrified directly on cryo-EM grids without any additional manipulation. We resolved a small but distinct population of hybrid lattice in both VLP preparations. In both cases, hybrid regions appeared as localized patches within the broader lattice network, often adjacent to immature or mature areas, suggesting that these sites may represent areas where structural remodeling is initiated or in progress during maturation (**Fig. 2A**). In the CA-NC VLPs, final particle counts indicated that 2.3% of regions adopted the hybrid lattice (**Fig. 2B, 2C**), whereas the remaining 97.7% displayed the immature lattice. Consistently, the CA-SP1 VLPs contained approximately 2.9% hybrid lattice, with the remaining representing a mixture of immature (41.5%) and mature (55.6%) lattices (**Fig. 2C**). Because the CA-SP1 VLPs are a naturally occurring maturation intermediate that display immature, hybrid, and mature features within the same VLP, this observation supports that the hybrid lattice represents a bona fide maturation transition state. Its presence in native, cell-derived VLPs indicates that this lattice state can arise intrinsically during the structural remodeling of Gag.

**Figure 2.**
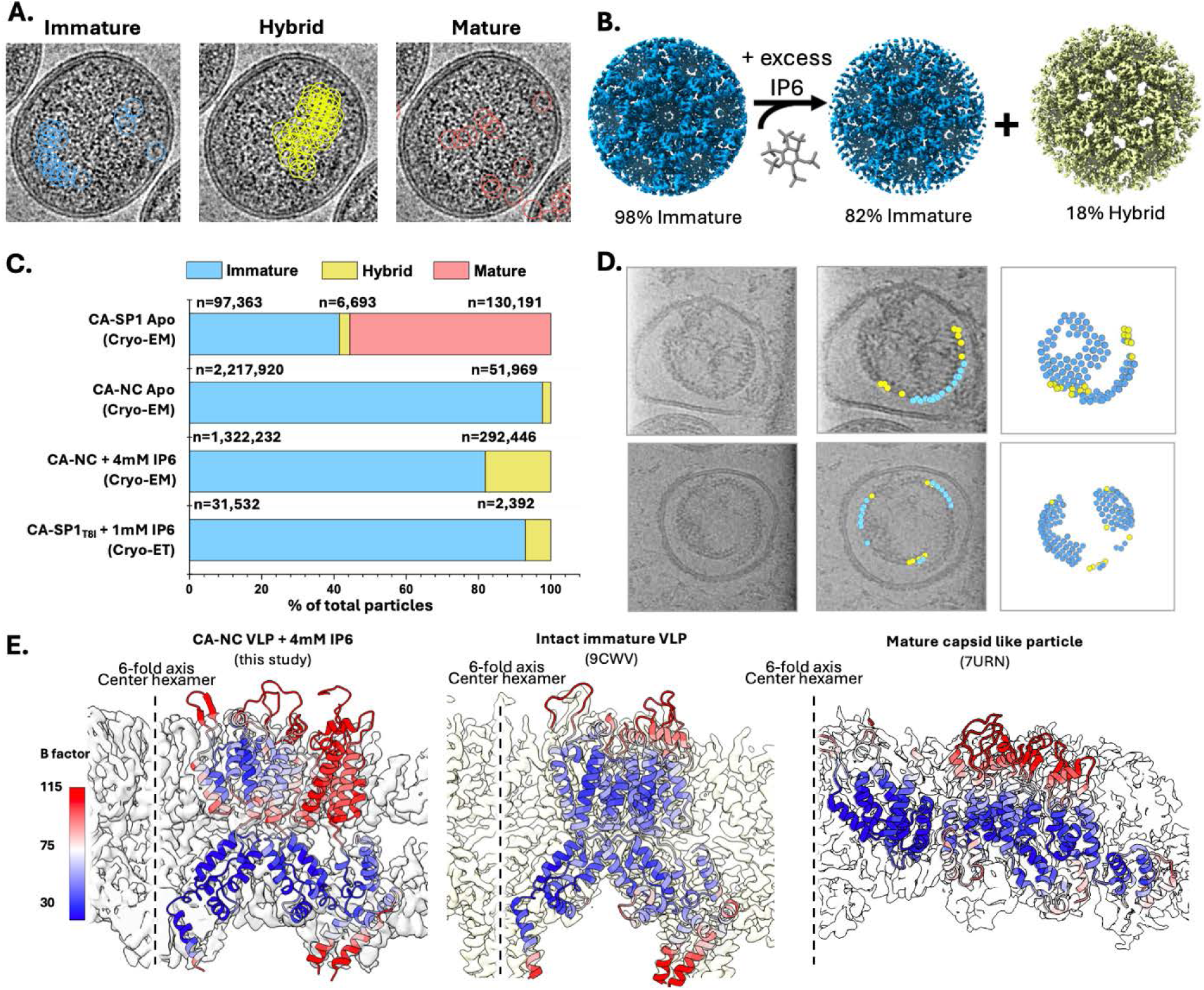
Hybrid lattice formation and spatial distribution in HIV-1 VLPs. A. Representative micrographs showing the location of immature (left, blue), hybrid (center, yellow), and mature (right, salmon) particles within the same CA-SP1 VLP. B. IP6-facilitated transition of lattice states. The immature lattice cryo-EM maps from CA-NC VLPs with (middle) or without (left) exogenous IP6 treatment are shown in blue, and hybrid lattice cryo-EM map from CA-NC VLPs treated with IP6 is shown in yellow. Treatment with excess IP6 enriched the hybrid lattice from 2% to 18%. Note the large, triangular shaped openings in the hybrid lattice. C. Bar graph showing the percentage of VLPs adopting the immature (blue), hybrid (yellow), and mature (salmon) lattice conformations as determined from cryo-EM and cryo-ET datasets. Percentages were calculated from the total number of particles in the dataset. *n* represents the total number of particles classified for each lattice type. D. Left, representative cryo-electron tomogram slices of individual HIV-1 CA-SP1_T8I_ VLPs treated with 1 mM IP6. Middle, spatial mapping of lattice types onto the corresponding tomogram slices in the left panel, with hybrid lattice segments colored yellow and immature lattice segments colored blue. Right, three-dimensional surface renderings of the mapped lattices within the same VLP. Hybrid lattices are consistently localized near points of lattice discontinuity. Snapshots are shown from different viewing angles corresponding to supplementary movies 1 and 2. E. B factor analyses of hybrid lattice from CA-NC VLPs treated with exogenous IP6 (left), immature VLPs (9CWV) (middle), and mature capsid-like particles (7URN) (right).

Given the presence of the hybrid lattice within cell-produced CA-NC and CA-SP1 VLPs, we next examined factors contributing to its formation. One key factor is IP6, a known regulator of HIV-1 assembly and maturation. Our structure of the hybrid lattice revealed that it incorporated three IP6 molecules—two occupying the canonical mature CA_NTD_ binding sites (**Fig. 1E and S3A**) and one positioned in the canonical immature CA_CTD_-SP1 binding site (**Fig. 1E and S3B**). This suggests that IP6 binding may promote the hybrid lattice by facilitating the transition of CA_NTD_ toward a more mature-like conformation from the immature form. To test this hypothesis, we used perfringolysin O (PFO) to permeabilize the VLP membrane and introduce additional exogenous IP6. Upon the addition of exogenous IP6 to CA-NC VLPs, the hybrid lattice population was markedly enriched, with the hybrid particle fraction increasing from 2.3% to 18.1% (**Fig. 2B, 2C**). The latter findings highlight the critical role of IP6 in stabilizing the hybrid lattice by promoting a conformational shift in CA_NTD_, reinforcing its function as a key regulator of HIV-1 maturation states. The shift from a predominantly immature lattice to a mixed population of immature and hybrid lattices upon excess IP6 addition underscores the dynamic nature of the hybrid lattice, consistent with its role as a transition state and with IP6 facilitating the structural progression toward this lattice form.

From our structural data, the transition from the immature to the hybrid and ultimately to the mature CA lattice necessitates substantial structural rearrangements within the virus particle. In a continuous pathway, CA_NTD_ rotates into the hybrid conformation, followed by lattice expansion as CA_CTD_ shifts toward CA_NTD_ to form mature interhexamer contacts. These conformational changes, particularly the large lattice expansion required, suggest that partial lattice disassembly is necessary to accommodate this transition.

Our single-particle cryo-EM analysis identified immature, hybrid, and mature CA lattices across the VLP population, but could not resolve how these distinct architectures are spatially organized within individual particles. To address this, we performed cryo-ET with subtomogram averaging of CA-SP1_T8I_ VLPs treated with PFO and exogenous IP6 to determine the structural arrangement of the lattice types. From the reconstructed tomograms, immature and hybrid lattices were refined to resolutions of 6.4 Å and 10.6 Å, based on 31,532 and 2,392 particles, respectively. Notably, when these particles were mapped back onto the original tomograms, we consistently observed both conformations within the same VLP (**Fig. 2C, 2D**, **and Supplemental Movie 1 and 2**). Furthermore, hybrid lattice regions were frequently localized near lattice discontinuities, suggesting that the transition occurs preferentially at the lattice edges, potentially facilitated by local disassembly during structural rearrangement (**Fig. 2D and Supplemental Movie 1 and 2**).

CA_NTD_ of the hybrid lattice exhibits elevated B factors, suggesting greater flexibility than CA_CTD_ than in the other lattice forms (**Fig. 2E**). Analyzing the *in situ* hybrid lattice model from CA-NC VLPs, the average B factor of CA_NTD_ was calculated as 88 Å^2^, which is significantly higher than those of CA_CTD_ (65 Å^2^), or CA_NTD_ from the immature Gag lattice (73 Å^2^, **Fig. 2E, middle**) and the mature capsid (69 Å^2^, **Fig. 2E, right**). When comparing the individual CA-SP1 chains from the center to the edge of the reconstruction, we observed a rapid rise of B factor for chains toward the edge (**Fig. 2E**). Although cryo-EM alignment typically incurs greater positional and orientational uncertainty near the edges of the processing box, reflected as elevated B factors, the hybrid lattice CA_NTD_ shows a steeper increase in B factor compared to immature and mature CA_NTD_. This observation suggests increased motion of hybrid CA_NTD_ and looser inter-subunit interactions, consistent with a potential transitional state. Furthermore, it aligns with our finding that hybrid lattice regions preferentially form at the periphery of the immature Gag lattice and near sites of lattice discontinuity, locations that are inherently less stable.

The localization of hybrid regions near gaps in the immature lattice supports a model in which partial disassembly of the immature lattice at its edges facilitates a displacive transition into the hybrid conformation. This spatial organization could also accommodate subsequent local expansion during the conformational rearrangements required for maturation. Furthermore, because additional IP6 molecules stabilize the hybrid lattice, disassembly of neighboring immature regions could increase local IP6 availability and thereby favor these transitions. Together, our observations suggest that both de novo reassembly and displacive remodeling may contribute to maturation, providing a flexible and spatially coordinated mode of lattice reorganization. These findings further raise the possibility that the hybrid lattice could serve as a nucleation site for the reassembly of disassembled capsomers.

### Comparison of the hybrid lattice with the immature and mature lattices

To further dissect the molecular features of the hybrid lattice observed *in situ*, we generated an *in vitro* CA-SP1 assembly system that could capture this intermediate state under controlled conditions and allow high-resolution structural analysis. Using the liposome-templating approach to promote efficient lattice formation^18, 33, 34^, we found that stabilizing the CA_CTD_-SP1 six-helix bundle was required to recapitulate the hybrid lattice observed *in situ*. In the absence of immature-state stabilization, wild-type CA-SP1 preferentially assembles into a mature CA lattice, consistent with CA_CTD_-SP1 dynamically interconverting between transiently disordered and stabilized six-helix bundle states and refolding to accommodate the mature capsid architecture^35^. Because the hybrid lattice adopts an immature-like CA_CTD_ arrangement, stabilizing the immature six-helix bundle was critical for trapping this intermediate. Accordingly, the addition of the maturation inhibitor BVM or incorporation of the six-helix bundle-stabilizing SP1_T8I_ mutation yielded assemblies comprised entirely of hybrid lattice (**Fig. S1**). These findings indicate that the hybrid lattice is unlikely to accumulate under unstabilized conditions, consistent with a rapid and coordinated immature-to-mature lattice transition, and provide an explanation for why this intermediate has not been observed previously. This stabilized *in vitro* reconstitution, therefore, enables targeted trapping of the hybrid state for mechanistic studies.

The newly observed hybrid lattice adopts a chimeric architecture comprising two layers: a CA_NTD_ layer in a mature-like conformation and a CA_CTD_-SP1 layer matching that of the immature Gag hexamer (**Fig. 1D**). This lattice architecture arises from a distinct arrangement between the CA_NTD_ and CA_CTD_-SP1 regions, with individual domains remaining essentially unchanged relative to the mature CA_NTD_ and immature CA_CTD_-SP1 structures (RMSD of 0.7 Å and 0.4 Å, respectively) (**Fig. S2**). The structural differences between the hybrid, immature, and mature lattices are enabled by flexibility in the interdomain linker, which permits reorientation between the CA_NTD_ and CA_CTD_. The primary difference stems from distinct hinge conformations involving residues M144 to I150 (**Fig. 3A**). The hybrid hinge adopts an intermediate geometry between the immature and mature states, enabling the hybrid relative orientation between CA_NTD_ and CA_CTD_ (**Fig. 3A**). Consistent with this, aligning a hybrid CA_CTD_ with an immature CA_CTD_ reveals a ∼140 degrees rotation of the hybrid CA_NTD_ relative to the immature CA_NTD_, whereas aligning a hybrid CA_NTD_ with a mature CA CA_NTD_ yields a ∼45 degrees reorientation in the hybrid CA_CTD_ relative to the mature CA_CTD_ (**Fig. 3B**).

**Figure 3.**
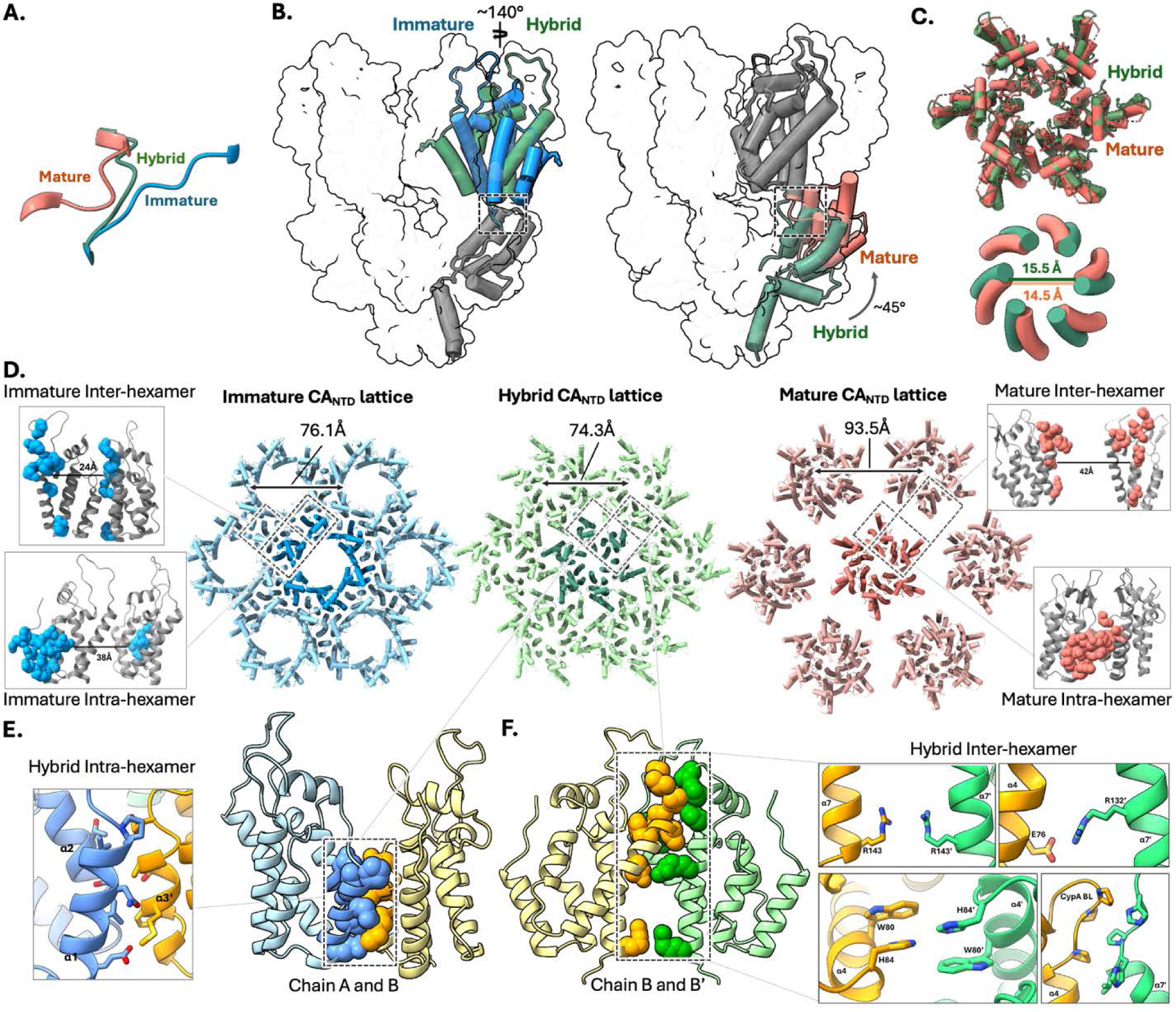
Structural comparisons of immature, hybrid, and mature lattices. A. Comparison of the distinct conformation of the CA NTD-CTD linker region (residues M144–I150) across the three structures (immature in blue, hybrid in green, and mature in salmon). B. Structural alignment of a CA monomer in the hybrid lattice (green) with that in the immature lattice (blue) (left) and with that in the mature lattice (salmon) (right). The flattened cryo-EM map of the hybrid lattice is displayed as background. The aligned domains are in grey, whereas the divergent domains are in their respective colors. CA_CTD_–aligned comparison shows the hybrid CA_NTD_ is rotated by ∼140 degrees relative to the immature CA_NTD,_ whereas CA_NTD_–aligned comparison shows the hybrid CA_CTD_ is rotated by ∼45 degrees relative to the mature CA_CTD_. Residue R18 is shown in sticks. Dashed box marks the region enlarged in panel A. C. Comparison of hybrid and mature CA_NTD_ hexamers. Top, superposition of the two CA_NTD_ hexamers. Bottom, measurement of the central pore diameter in CA_NTD_ hexamers reveals a pore of 15.5 Å in the hybrid hexamer versus 14.5 Å in the mature hexamer. D. Top-down view of the CA_NTD_ layer for seven neighboring hexamers in the immature (blue, center left), hybrid (green, center middle), and mature (salmon, center right) lattices, with the central hexamer in a darker shade. The hybrid lattice CA_NTD_ is closely packed, whereas the mature lattice shows increased spacing with no direct inter-hexamer CA_NTD_ contacts. Distances shown are between R18 Cɑ atoms. Dashed boxes mark three monomer chains illustrating distinct lattice interfaces. The figure insets show that CA_NTD_ interface residues in the hybrid lattice (spheres) are separated in the corresponding immature inter-hexamer (top left), immature intra-hexamer (bottom left), and mature inter-hexamer (top right) interfaces. Only the mature intra-hexamer interface (bottom right) preserves a similar contact. E. Distinct interactions at the hybrid CA_NTD_ intra-hexamer interface between chains A (blue) and B (orange), which are absent in the immature lattice. The left inset shows a close-up view of the interface. F. Distinct interactions at the hybrid CA_NTD_ inter-hexamer interface between chains B (orange) and B’ (green), which are absent in the immature and mature lattices. The right insets show close-up views of selected interacting residues, including those involving the CypA-binding loop (CypA BL’, bottom right).

At the assembly level, the CA_CTD_-SP1 layer of the hybrid lattice is essentially unchanged from that of the immature Gag lattice (RMSD of 0.5 Å for the hexamer). In contrast, the CA_NTD_ layer of the hybrid lattice exhibits distinct features compared to the mature form. First, the hybrid and mature CA_NTD_ hexamers are broadly similar with notable deviations (RMSD of 3.8 Å for the hexamer), and the hybrid hexamer shows a slightly more open central pore (**Fig. 3C**). Second, and more prominently, the CA_NTD_ hexamers pack more densely in the hybrid lattice and make direct inter-hexamer contacts, whereas the mature CA_NTD_ hexamers are spaced farther apart and lack direct contacts (**Fig. 3D**), with lattice instead mediated by CA_CTD_ interactions. Quantitatively, the inter-hexamer spacing, measured as the Cα–Cα distance between R18 residues of neighboring hexamers, is 76.1 Å in the immature lattice, 74.3 Å in the hybrid lattice, and 93.5 Å in the mature lattice, highlighting the substantially expanded packing of the mature hexamers relative to the immature and hybrid states (**Fig. 3D**). Together, these observations support a maturation-associated expansion from tightly packed hexamers (immature and hybrid) to a markedly more open mature lattice, a transition that likely entails local lattice disruption near hybrid transition sites—consistent with their enrichment near immature lattice discontinuities, where partial disassembly and reassembly can be most readily accommodated.

### Interface analysis of the hybrid lattice

The newly identified hybrid lattice contains a CA_CTD_-SP1 layer indistinguishable from that of the immature Gag_CA-SP1_ structure. In contrast, the mature-like hybrid CA_NTD_ forms distinct inter-hexamer interfaces, whose structural details are resolved in our high-resolution cryo-EM map. To describe the two principal interface classes, we denote adjacent CA chains within the same hexamer as A and B, and a CA chain from a neighboring hexamer as B’, defining the intra-hexamer interface as A:B and the inter-hexamer interface as B:B’ (**Fig. 3E**, **3F**). Notably, the inter-hexamer spacing produces large, triangular openings that create open channels that may facilitate solvent accessibility.

The intra-hexamer A:B interface buries a surface area of 1,144 Å², comprising the canonical CA_CTD_-SP1:CA_CTD_-SP1 interaction as observed in the immature Gag_CA-SP1_ lattice together with a slightly altered mature-like CA_NTD_:CA_NTD_ interface. This interface is mediated primarily by an α-helix (residues 38–45) from chain A and α-helices (residues 15–28 and 54–60) from chain B, with contacts largely driven by hydrophobic interactions (**Fig. 3E**). The hybrid intra-hexamer CA_NTD_ interface differs from that of the immature lattice but closely resembles the mature lattice (**Fig. 3D**), underscoring the mature-like organization of the CA_NTD_ in the hybrid state.

The inter-hexamer B:B’ interface is unique to the hybrid lattice and buries 1,068 Å² of surface area (**Fig. 3F**). The contacts are distributed across the CA_NTD_, interdomain hinge, and CA_CTD_-SP1 regions. The CA_CTD_-SP1:CA_CTD_-SP1 interface again matches that in the immature Gag_CA-SP1_ lattice, whereas the CA_NTD_ interfaces are distinct from both the immature and mature lattices (**Fig. 3D**). At the interdomain hinge, two arginine residues (R143) from chains B and B’ engage in a π-stacking interaction (**Fig. 3F**). The inter-hexamer CA_NTD_:CA_NTD_ interface bridges adjacent hexamers and is absent in the mature CA lattice, where CA_NTD_ regions from neighboring capsomeres are distant and do not interact (**Fig. 3D**). In the more compact hybrid lattice, this interface is extensive and stabilized by a network of interactions (**Fig. 3F**): (1) a salt bridge between E76 and R132’; (2) symmetrical interactions between W80 and H84 from each chain; and (3) hydrophobic interactions between V83-H87 in the Cyclophilin A binding loop and P122’-P125’ in a proline-rich loop.

The large separation between chains A and B’ defines a prominent solvent-accessible channel within the hybrid lattice. From a top-down view, this region adopts a distinctive triangular geometry that serves as a structural hallmark of this lattice type. Although residues that contribute to the FG-binding pocket in the mature capsid, including N74, K70, and N57, are oriented toward this channel (**Fig. S4A and S4B**), the local architecture differs substantially from that of a fully formed binding site. In particular, the hybrid lattice lacks the neighboring CA_CTD_ interactions that cap and stabilize the pocket in the mature lattice (**Fig. S4C**), a feature known to be critical for high-affinity LEN binding^34–37^. Consistent with this interpretation, LEN treatment shifted the population from a uniform hybrid lattice toward a mixture of hybrid and mature CA lattices (**Fig. S4D**), yet no corresponding LEN density was observed in the remaining hybrid particles. This suggests that LEN does not stably bind the hybrid lattice, but instead preferentially stabilizes mature-like configurations once appropriate CA_CTD_ contacts are established. Together, these observations support a model in which the hybrid lattice represents a structurally permissive intermediate that can be driven toward maturation by LEN, despite lacking the fully assembled binding site required for direct, stable inhibitor engagement.

### Mutations that specifically destabilize the hybrid lattice prevent proper virus maturation

To validate that the hybrid lattice is required for HIV-1 maturation, we sought to selectively destabilize the hybrid lattice without disrupting immature lattice formation. We hypothesized that if the hybrid lattice is a necessary intermediate in the maturation pathway, then impairing its formation would prevent proper maturation even if immature virion assembly and budding remained intact.

Based on the hybrid-specific interface (**Fig. 3F**), we generated a mutation designed to selectively disrupt the hybrid CA_NTD_ inter-hexamer interface and assessed its effect on virus production and maturation (**Fig. 4A**). H84 is well-suited for this test because it participates in the hybrid interface but not in any immature or mature lattice interfaces. The H84A mutant produced virus particles, with modestly reduced p24 production relative to the R9 wild type (WT) control (**Fig. 4B**), indicating that the immature Gag lattice can still assemble and facilitate virion budding in the presence of the mutation. In contrast, the released mutant virions were non-infectious (**Fig. 4C**), consistent with prior observations that this substitution produces assembly-competent but poorly infectious virions^36^. To more directly probe capsid integrity, we compared natural endogenous reverse transcription (NERT) and endogenous reverse transcription (ERT): both assays measure intravirion DNA synthesis, but ERT is performed after membrane permeabilization such that efficient reverse transcription requires a well-assembled capsid to maintain an enclosed reaction environment, whereas NERT is carried out in intact virions and thus serves as a capsid-independent control for enzymatic competence. Consistent with this rationale, H84A retained robust NERT activity but exhibited markedly reduced ERT activity, resulting in a strongly decreased ERT/NERT ratio for both minus-strand strong-stop (MSS) and full-length minus-strand (FLM) DNA products (Fig. 4D). Together, these data indicate that although H84 does not mediate a direct contact in the mature capsid lattice, its perturbation nonetheless compromises mature core integrity, consistent with an indirect role in maturation— most plausibly by disrupting a transient intermediate, such as the hybrid lattice, that is required to progress to the mature architecture.

**Figure 4.**
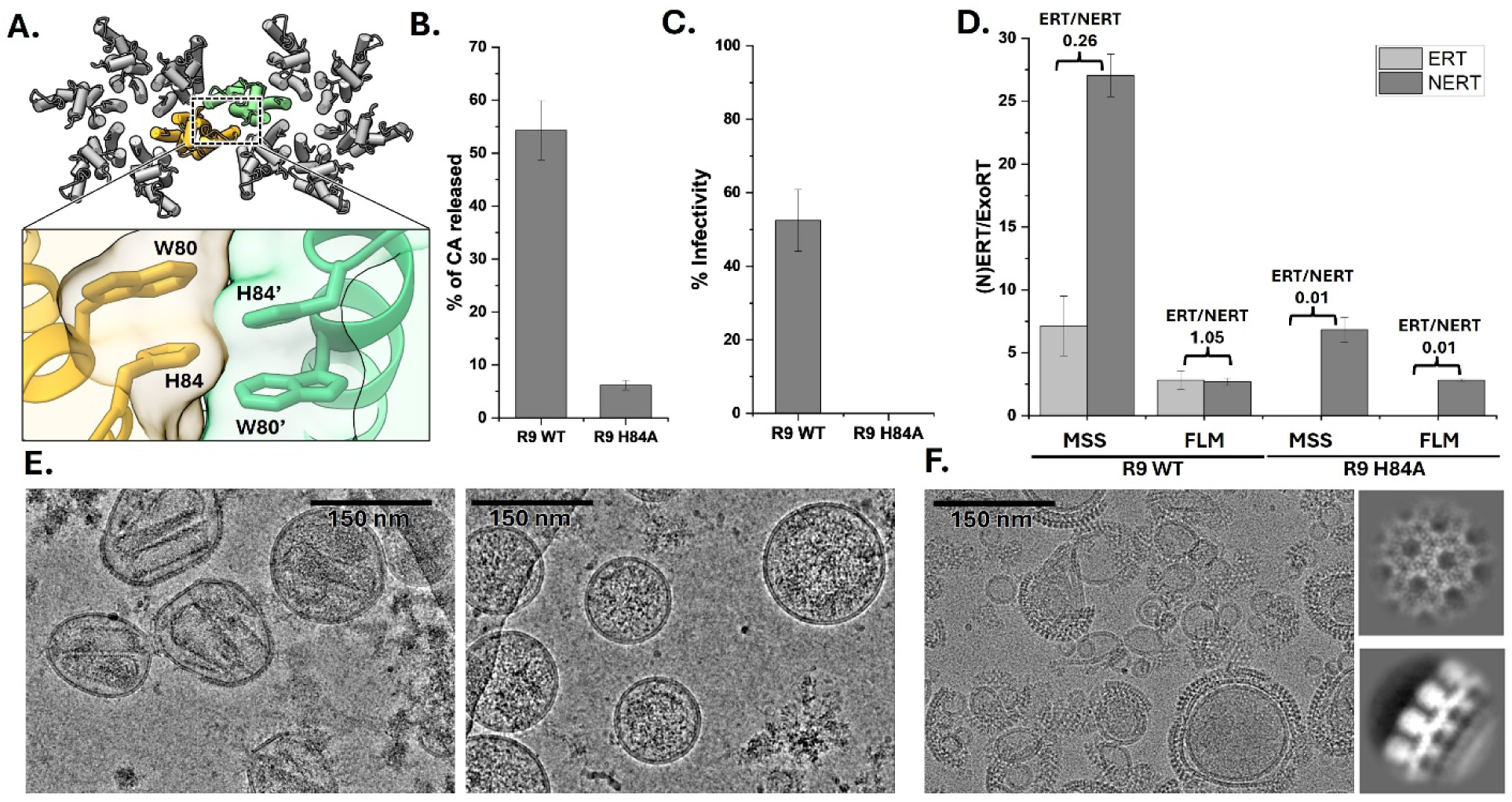
H84A mutation impairs HIV-1 maturation and infectivity. A. Illustration highlighting the locations and interactions of the neighboring H84 residues within the inter-hexamer interface of the hybrid lattice. B. p24 production from R9 WT and R9 H84A VLPs. Error bars indicate the standard error of the mean from triplicate experiments. C. Bar graph showing the infectivity of R9 H84A VLPs compared to R9 WT VLPs, normalized by ExoRT activity. Error bars indicate the standard error of the mean from triplicate experiments. D. Bar graph showing the ERT and NERT activity, normalized by ExoRT, for R9 WT and R9 H84A viruses. MSS (minus-strand strong-stop DNA) products and FLM (full-length minus-strand DNA) products are shown for each sample. ERT products are shown in light grey and NERT products are shown in dark grey. ERT/NERT ratios are shown above each sample. Error bars indicate the standard error of the mean from triplicate experiments. E. Cryo-EM images of R9 WT (left) and R9 H84A (right) VLPs highlighting the absence of a matured core in the H84A mutant sample. F. H84A CA-SP1 assemblies on SUV liposomes; right panels show 2D class averages confirming an immature lattice arrangement.

To further validate the structural consequences of H84A, we examined VLPs with this mutation by cryo-EM and found no evidence of a mature CA lattice (**Fig. 4E**), consistent with a failure to form an appropriate mature core. Despite impaired maturation, we hypothesized that immature lattice assembly can still proceed, given that virion release still occurred in the H84A background. We tested the hypothesis by introducing H84A into our *in vitro* immature Gag assembly system. As expected from the lack of H84 involvement in immature lattice interfaces, the mutant efficiently assembled on liposomes and formed an immature lattice comparable to WT (**Fig. 4F**). These results confirm that the hybrid-specific H84A mutation permits immature virus assembly but prevents progression to the mature state.

Together, these results show that a hybrid-destabilizing substitution can preserve immature assembly and particle release yet compromise the downstream lattice remodeling required for maturation and infectivity. These observations support a maturation pathway in which HIV-1 Gag lattice reorganization proceeds, at least in part, through a displacive process that generates a transient hybrid lattice. In this framework, the hybrid lattice could function as an intermediate and a local nucleation point for reassembly of disassembled capsomers during maturation.

### CA NTD-CTD linker dynamics and neighboring loop remodeling coordinate conformational transitions during maturation

We used targeted molecular dynamics (TMD) simulations to further probe the structural maturation of Gag and elucidate how the hybrid lattice bridges the maturation process. These simulations describe conformational rearrangements at the monomer level and modeled transition pathways linking the immature, hybrid, and mature states, revealing how biochemical cleavage events drive stepwise structural remodeling of the lattice to form the mature capsid. To comprehensively explore the conformational landscape, the TMD trajectories were guided between the states in both clockwise and counterclockwise directions, allowing us to assess whether multiple transition routes could contribute to lattice morphing during maturation (**Fig. S5**). The structural transitions between the immature and hybrid states were evaluated using quaternion-based 3D rotations, whereas those between the hybrid and mature states were evaluated using conventional RMSD-driven translations (**Fig. S5**).

A prominent feature emerging from these simulations is the central role of the CA NTD-CTD linker in mediating the transition between lattice states during maturation. Our TMD simulations indicate that the structural rearrangement occurs progressively during clockwise rotation, as reflected in the Ramachandran plot analysis (**Fig. 5)**. In the immature conformation (**Fig. 5A**, left), Y145 occupies a tightly packed position wedged between P147 and residues F32 and H62, maintaining an extended backbone. Upon transition to the hybrid conformation (**Fig. 5A**, middle), Y145 rotates outward toward the solvent, shifting the backbone into a more helical configuration (**Fig. 5B**). As hybrid to mature transition proceeds, the P145 backbone first adopts a helical configuration, then straightens, and finally forms a helical turn as the CA_CTD_ is positioned, with P147 adjusting in concert to stabilize the new arrangement (**Fig. 5C**). Furthermore, once Y145 exits the packed geometry of the immature conformation, its sidechain remains relatively flexible until it ultimately locks into the mature orientation (**Fig. 5A**). Together, these coordinated residue-level rearrangements illustrate how local structural transitions propagate through the linker to support the larger-scale lattice remodeling required for maturation.

**Figure 5.**
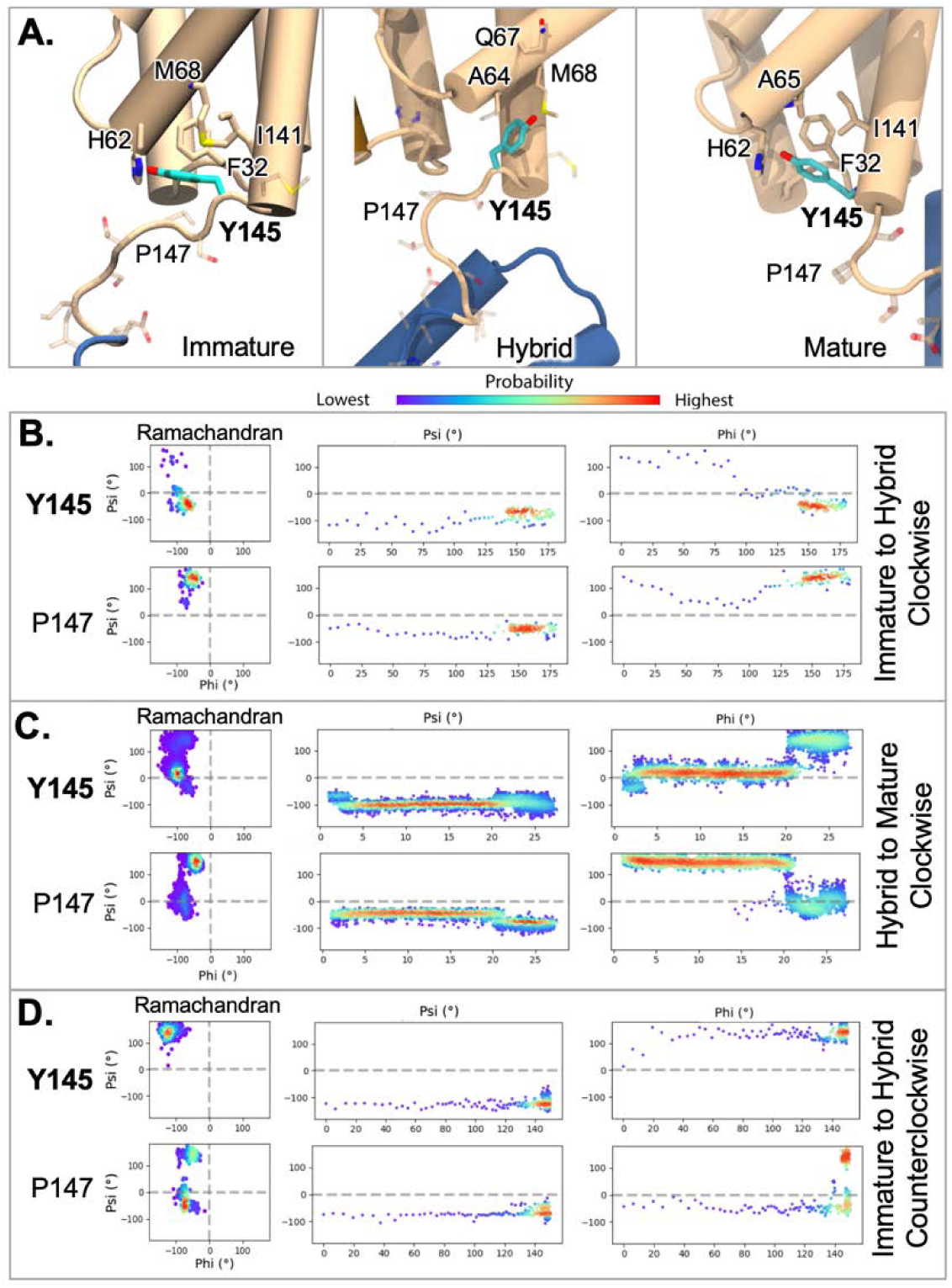
The CA NTD-CTD linker rearrangements during structural maturation. A. Snapshots of the orientation of Y145 in the immature, hybrid, and mature monomers, with the surrounding residues (sticks) within 5Å, illustrate the differences in the Y145 packing. B. Ramachandran plots for Y145 and P147 indicating two-state behaviors in the clockwise immature-to-hybrid pathway, with the trace plots for the dihedral angles (middle and right). C. Ramachandran plots for Y145 and P147 indicating two-state behaviors in the clockwise hybrid to mature transition, with the trace plots for the dihedral angles (middle and right). D. Ramachandran plots for Y145 and P147 indicating two-state behaviors in the counterclockwise pathway, with the trace plots for the dihedral angles (middle and right).

A second region that undergoes coordinated rearrangement is the loop (residues 174-178) between helices H8 and H9 in CA_CTD_ (**Fig. S6**). In the immature state, Q176 contacts E28 at the C-terminus of H1 in the CA_NTD_, an interaction that is lost as the rotation toward the hybrid state begins. During the transition under lower force constants, Q176 transiently engages residues in the CA_NTD_ H3/H4 loop (H62 and Q63), suggesting an intermediate stabilization step as the linker progresses from the immature to hybrid state. Under higher force constants, however, these contacts are absent, and the faster rotational progression likewise limits proper adoption of the hybrid configuration of the CA NTD-CTD linker. Together, these observations indicate that coordinated interactions between the H8/H9 and H3/H4 loops help steer the linker into the hybrid arrangement and that this effect is sensitive to the kinetics of the rotational transition.

In the counterclockwise reverse (hybrid-to-immature) rotation, the pathway fails to restore the tightly packed Y145 configuration in the immature form, even under strong energetic bias (**Fig. 5D**). The simulations show that once this packing is disrupted, it does not readily reestablish, suggesting that hybrid formation is directionally favored and may represent a committed step toward the mature state. P147 undergoes a coordinated reorganization with Y145, further supporting the directional nature of this transition. This inability to return to the immature conformation arises also from steric interference during the reverse rotation, during which the CA_NTD_ clashes with residues E29 and K30 of a neighboring subunit. In the mature lattice, residues E28, E29, and K30 line the central channel, while Q176 contacts R143 at the CA NTD–CTD interface^37–39^, collectively biasing the transition toward forward progression.

Together, these results support a model in which the hybrid conformation acts as a bona fide intermediate in the HIV-1 maturation pathway rather than an off-pathway state. The coordinated reorganization of Y145, P147, and Q176 illustrates how CA_NTD_ rotation and linker repositioning generate stabilizing interactions that drive the lattice toward forward progression. Prior studies show that Y145A substitution produces non-infectious virions with unstable, aberrant cores, highlighting how perturbation of this linker region disrupts proper capsid assembly and stability^40, 41^. Importantly, the inability of the system to restore the tightly packed immature Y145 configuration, even under strong reverse bias, suggests an inherent directionality imposed by a ratchet-like mechanism.

## Discussion

HIV-1 maturation has been framed by three mechanistic scenarios: a displacive pathway in which the immature lattice is remodeled into the mature lattice through coordinated structural rearrangements^22, 23^, a de novo pathway in which the immature lattice disassembles and the mature lattice reassembles^19–21^, and hybrid schemes that combine both processes^25^. Despite extensive study, structural insight into how these two lattice endpoints are connected has remained limited. Here, we identify an additional lattice architecture that is distinct from both the immature and mature forms and combines an immature-like CA_CTD_-SP1 organization with a mature-like CA_NTD_ fold. This intermediate lattice provides a concrete structural linkage between the two endpoint architectures and suggests a route by which maturation could proceed through localized displacive remodeling events, potentially coordinated with partial disassembly and reassembly.

Our data show that the hybrid lattice occurs not only *in vitro* but also in native, enveloped CA-NC and CA-SP1 VLPs produced from human cells^42, 43^, demonstrating that this conformation can arise under physiological conditions. Building on this, we find that IP6 also modulates hybrid lattice formation, extending its established role in HIV-1 lattice organization and structural transitions^11–16^. The hybrid lattice incorporates three IP6 molecules—two at the mature CA_NTD_ binding sites and one at the immature CA_CTD_ site—providing a mechanistic basis for how IP6 can stabilize this intermediate architecture. Consistently, while CA-NC VLPs primarily assemble into the immature lattice under standard conditions, the addition of excess IP6 promotes the formation of a mixed population, enriching the hybrid lattice. These observations extend the established role of IP6 in lattice assembly by implicating it in tuning the structural balance between immature, hybrid, and mature lattice states during maturation.

Our combined structural and mutational analyses support a model in which the hybrid lattice represents an intermediate in HIV-1 CA lattice remodeling. Cryo-EM and cryo-ET of VLPs revealed the coexistence of immature, hybrid, and mature lattices within the same particle, with hybrid regions often enriched near lattice discontinuities. This spatial organization suggests that conversion toward the hybrid lattice may start at immature lattice edges, where reduced lattice continuity can accommodate local conformational rearrangements and partial disassembly. Edge-associated disassembly would also release IP6 to increase its local availability, which could further bias the lattice toward the IP6-stabilized states. In this context, the additional degrees of freedom at lattice boundaries can accommodate the CA_NTD_ rotation characteristic of the hybrid lattice and the subsequent lattice expansion associated with CA_CTD_ reorientation required for maturation. Together, these results are consistent with a maturation process that is spatially and temporally coordinated and involves displacive remodeling coupled to localized disassembly and reassembly.

A mutation designed to specifically disrupt a key interface unique to the hybrid conformation further supports a functional role for this architecture during maturation. The H84A substitution did not prevent immature lattice formation or virus budding, yet the resulting particles were non-infectious, showed a loss of ERT activity consistent with defective mature core formation, and displayed severely disrupted capsid morphology. Importantly, H84A supported immature lattice assembly both in cells and *in vitro*, indicating that the defect arises during downstream lattice remodeling rather than during Gag assembly. Moreover, because H84 does not participate in mature lattice interfaces and is not required for the mature CA fold, the phenotype is most consistent with disruption of an upstream remodeling step en route to the mature lattice. Taken together, these results lend strong support to the hypothesis that the hybrid lattice is an important intermediate step between the immature and mature states (**Fig. 6**).

**Figure 6.**
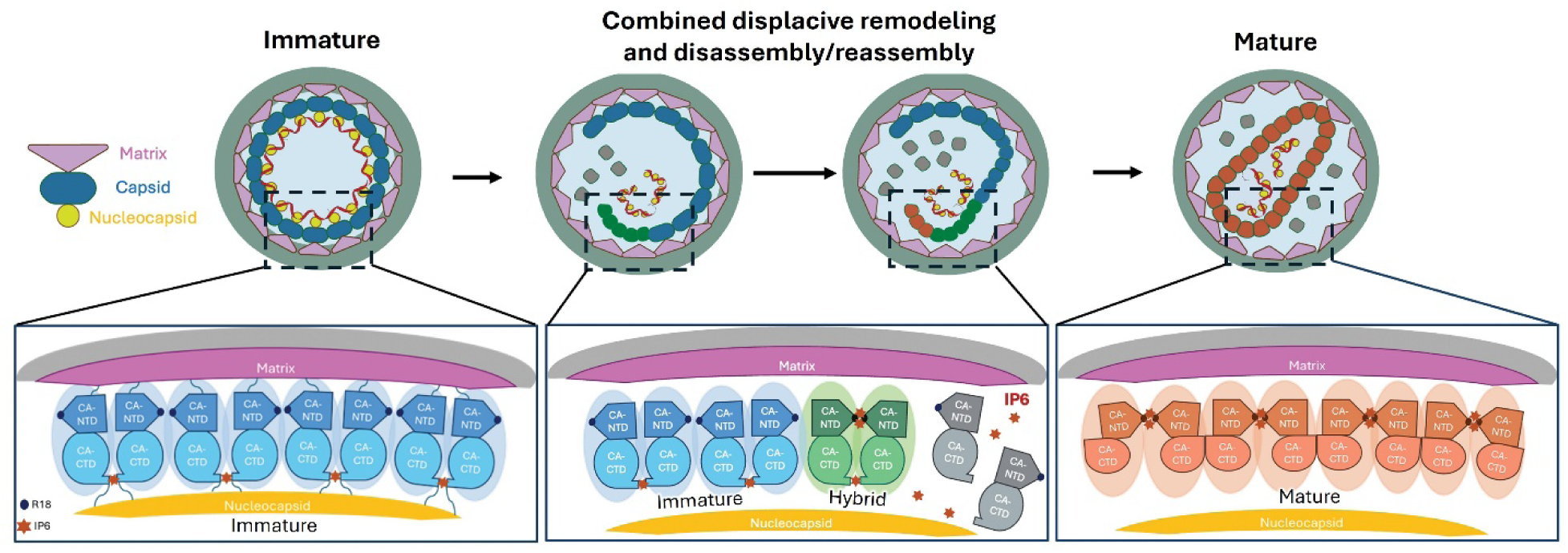
HIV-1 Maturation Pathway Highlighting the Hybrid Lattice Transition. Model of HIV-1 maturation showing the immature lattice (left) transitioning to the mature lattice (right). During the intermediate stage (center), some capsomeres remain immature, others disassemble, and some adopt the hybrid conformation. In the hybrid lattice, CA_NTD_ rotates and binds two IP6 molecules per hexamer while CA_CTD_-SP1 retains its immature state bound to one IP6. Subsequent CA_CTD_ rearrangements drive formation of the mature, conical capsid (right). Capsomeres are colored to indicate their conformational states: immature (blue), hybrid (green), mature (salmon), and disassembled (grey).

Our TMD simulation results suggest that HIV-1 capsid maturation is driven by coordinated rearrangements centered on the CA NTD-CTD linker and neighboring loops. Specifically, Y145 and P147 undergo coupled shifts that reposition the linker, while the H8/H9 and H1/H2 loops form transient contacts that can guide domain rotation. In the simulations, release of Y145 from tight immature packing, together with steric constraints that arise during reverse CA_NTD_ rotation, bias the system away from the immature configuration and disfavor full reversal. Together, these features are consistent with a ratchet-like mechanism that channels immature lattice through the hybrid state toward the mature lattice, providing a plausible structural explanation for directionality during maturation.

In conclusion, by identifying and structurally defining a hybrid CA lattice that bridges immature and mature architectures, this study provides a previously missing structural framework for understanding how HIV-1 reorganizes its structural lattice during maturation. These results support stepwise remodeling in which localized displacive rearrangements are coupled to partial disassembly and reassembly (**Fig. 6**): disruption of portions of the immature lattice could increase local IP6 availability, enabling remaining CA-SP1 hexamers to bind IP6 and stabilize the hybrid lattice transition, which in turn may serve as a structural nucleus for subsequent lattice maturation as free CA or CA–SP1 subunits are reassembled. More broadly, this shifts maturation from a binary choice between displacive remodeling and de novo reassembly to a mechanistically testable continuum in which lattice plasticity, localized disassembly/reassembly, and small-molecule cofactors such as IP6 jointly shape the remodeling trajectory. This intermediate state also highlights a potential vulnerability in the maturation pathway, suggesting that hybrid lattice architectures—not only the immature assembly and mature capsid—may be tractable targets for therapeutic intervention. It also provides a conceptual framework for dissecting analogous remodeling processes in other retroviruses.

## Methods

### CA-SP1 and CA-NC protein preparations

The CA-SP1 (Gag 133-376) construct was cloned into the expression vector pET28b, incorporating a C-terminal TEV cleavage linker sequence “ENLYFQGGSS” and a tandem 6x His-tag. The protein was expressed in BL21 (DE3) *Escherichia coli*. The cells were cultured in Terrific Broth (TB) medium at 37°C, and expression was induced with 0.5 mM IPTG at 18°C once the OD600 reached 0.8. The cell pellets were resuspended in lysis buffer (50 mM Tris, pH 7.5, 0.5 mM TCEP, 300 mM NaCl) supplemented with 1X HALT protease inhibitor (Thermo Fisher Scientific) and lysed using a microfluidizer. Following centrifugation to remove cell debris, the soluble lysate was incubated with 35% ammonium sulfate and centrifuged to collect the protein precipitant. The precipitant was then solubilized and dialyzed in the buffer (50 mM Tris, pH 7.5, 0.5 mM TCEP, 300 mM NaCl). The solution was filtered through 0.45 µm filters and loaded onto a 5 mL HisTrap column (Cytiva) on an AKTA system (GE) at a flow rate of 1 mL/min. The column was washed with lysis buffer and eluted with a gradient of elution buffer containing an additional 500 mM imidazole. The final CA-SP1 sample was polished using size exclusion chromatography on a Superdex 200 column (GE), and the protein concentration was determined using a Nanodrop (Thermo Fisher Scientific). The final protein sample was concentrated to approximately 12 mg/mL, aliquoted, flash-frozen in liquid nitrogen, and stored at −80°C. The CA-SP1_T8I_ and CA_H84A_-SP1 constructs were generated using the Q5 Site-Directed Mutagenesis Kit (NEB) to generate the T8I and H84A point mutations. The same purification protocol described for the CA-SP1 construct was followed for the CA-SP1_T8I_ and CA_H84A_-SP1 constructs.

The CA-NC (Gag 133-432) construct was synthesized and cloned by Twist Biosciences into the expression vector pET28b. The protein was expressed in LOBSTR-BL21 (DE3) RIL *Escherichia coli* (Kerafast). The cells were cultured in TB medium at 37°C, and expression was induced with 1mM IPTG at 18°C once the OD600 reached 0.8. The cells pellets were resuspended in lysis buffer (50 mM Tris pH 7.5, 750 mM NaCl, 5% glycerol, 10 µM ZnCl_2_, 1 mM DTT, 0.5 mM TCEP) supplemented with 1X HALT protease inhibitor (Thermo Fisher Scientific) and lysed using a microfluidizer. Following centrifugation to remove cell debris, the soluble lysate underwent 0.4% PEI precipitation for 30 minutes at 4°C and was then subjected to centrifugation for 30 minutes at 21,000 x g to pellet nucleic acids. The supernatant was then treated with 30% ammonium sulfate for 2 hours at 4°C and was then subjected to centrifugation for 30 minutes at 21,000 x g. The pellet was resuspended in resuspension buffer (50 mM HEPES pH7.5, 50 mM NaCl, 5% glycerol, 10 µM ZnCl_2_, 0.5 mM TCEP, 1 mM DTT) and loaded onto a 5 mL HiTrap SP HP column (Cytiva) on an AKTA system (GE) at a flow rate of 1 mL/min. The column was washed with resuspension buffer and eluted with a gradient of elution buffer (50 mM HEPES pH7.5, 1 M NaCl, 5% glycerol, 10 µM ZnCl_2_, 0.5 mM TCEP, 1 mM DTT). The final CA-NC sample was polished using size exclusion chromatography on a Superdex 200 column (GE), and the protein concentration was determined using a Nanodrop (Thermo Fisher Scientific). The final protein sample was concentrated to approximately 16 mg/mL, aliquoted, flash frozen, and stored at −80°C for future use.

### Perfringolysin O (PFO) protein purification

The gene encoding the cytolytic toxin PFO from *Clostridium perfringens* was cloned into a pET-21-derived vector (Novagen) as a 6x His-tagged construct and expressed in BL21 (DE3) *Escherichia coli*. The cells were cultured in Terrific Broth (TB) media at 37°C with shaking until reaching an OD600 of 0.8. Protein expression was induced with 0.5 mM IPTG for 16 hours at 18°C, followed by cell collection via centrifugation. The cell pellets were stored at −80°C until use. For lysis, the pellets were resuspended in lysis buffer (50 mM Tris, pH 7.5, 0.5 mM TCEP, 300 mM NaCl) supplemented with 1X HALT protease inhibitor (Thermo Fisher Scientific) and lysed using a microfluidizer. After centrifugation to remove cell debris, the supernatant was filtered through 0.45 µm filters and loaded onto a 5 mL HisTrap column (GE) on an AKTA system (GE) at a flow rate of 1 mL/min. The column was washed with lysis buffer and eluted with a gradient of elution buffer containing an additional 500 mM imidazole. The purified PFO concentration was determined using a Nanodrop (Thermo Fisher Scientific). The final PFO sample, at a concentration of 10 mg/mL, had 15% glycerol added, was flash-frozen in liquid nitrogen, and stored at −80°C until use.

### Liposome preparation

The preparation method was adapted from Highland et al., 2023^18^. Stock solutions of 1,2-dioleoyl-sn-glycero-3-[(N-(5-amino-1-carboxypentyl)iminodiacetic acid)succinyl] nickel salt (DGS-NiNTA) and 1,2-dioleoyl-sn-glycero-3-phosphocholine (DOPC) in chloroform were obtained from Avanti Polar Lipids. Cholesterol was purchased from Thermo Scientific Chemicals and dissolved in chloroform at a concentration of 5 mg/mL. Lipids were combined in an 85:10:5 ratio of DOPC:DGS-NiNTA:Cholesterol using a volumetric flask, yielding a final mixture composed of approximately 95% DOPC, 2% DGS-NTA, and 3% cholesterol. This lipid mixture was transferred to a round-bottom flask and dried under rotary evaporation for 5 hours to create a thin lipid film. The dried lipids were then resuspended in buffer (25 mM HEPES pH 7.4, 150 mM KCl) to a final concentration of 13 mM by gentle agitation on a rotory evaporator. The suspension was left to hydrate overnight at room temperature. To produce large unilamellar vesicles (LUVs), the hydrated lipid suspension was extruded through a 100 nm polycarbonate membrane 100 times at 37°C.

### Protein assemblies

The CA-NC protein was assembled by combining the following components at the given final concentrations: 150 µM CA-NC protein, 1 mM IP6 (Sigma Aldrich), and 12 µM Ambion yeast tRNA (Roche) with 300 µM LEN (for the indicated sample containing LEN). The assembly mixture was then subjected to overnight dialysis at 4°C in the assembly buffer (25 mM HEPES, 50 mM NaCl, 10 µM ZnCl_2_, 5% glycerol, pH 8).

The CA-SP1 proteins were assembled by combining the following components at the given final concentrations: 150 µM CA-SP1 protein, 1 mM IP6 (Sigma Aldrich), 4 mM LUV, with 100 µM BVM (for the indicated sample containing BVM) in the assembly buffer (25 mM HEPES, 50 mM NaCl, 5% glycerol, pH 8). These samples were allowed to assemble for 1 hour at room temperature before subjecting them to further analysis.

### CA-NC and CA-SP1 VLP preparation

VLPs were expressed by transfecting 1 µg/mL CA-SP1 (CA5) or CA-SP1-NC (CA6, referred to as CA-NC in this study) DNA^42^ using the Expi293 Transfection Kit (Thermo Fisher Scientific) in Expi293F cells (Thermo Fisher Scientific), that were cultured in 8% CO_2_ at 37°C shaking at 250 rpm. 24 hours post transfection, Enhancer 1 and Enhancer 2 from the Expi293 Transfection Kit (Thermo Fisher Scientific) were added. 48 hours post transfection, the suspension of VLPs and cells were collected and centrifuged at 500 x g for 5 minutes to pellet the cells. The supernatant was then filtered through a 0.45 µm syringe filter. To isolate and purify the VLPs, the filtered supernatant was pelleted through at 20% sucrose cushion by ultracentrifugation in an SW32 Beckman rotor at 120,000 x g for 3 hours at 4°C, and the resulting pellet was resuspended in 600 µL 10 mM Tris-HCl, 100 mM NaCl, 1 mM EDTA, pH 7.4 (1X STE). VLPs resuspended in 1x STE buffer at a concentration of 3 mg/mL, were aliquoted, and frozen to −80°C for future cryo-EM sample grid preparation.

VLPs were prepared for structural analysis by incubating 100 µM VLPs with 1µM PFO for 10 min at 37°C with 4 mM IP6 for the experimental sample and no IP6 for the control sample. After 5-10 minutes, the sample was removed from 37°C and used for cryo-EM sample grid preparation immediately.

### Negative stain EM of samples

Glow-discharged (25 mA for 30 seconds) carbon-coated 300 mesh Cu grids (Electron Microscopy Sciences) was applied with 3.5µL of sample for 1 min. The grid was then washed with 2% uranyl acetate (Electron Microscopy Sciences), followed by staining in 2% uranyl acetate (Electron Microscopy Sciences) for 1 minute, blotted, and allowed to dry before subjected to imaging analysis. Imaging was performed on a 120 kV Talos L120C electron microscope (Thermo Fisher Scientific) with CETA CMOS camera.

### Cryo-EM data collection

Grids for cryo-EM were prepared by applying 4 µL of sample onto glow-discharged Quantifoil 300 mesh Cu 2/1 grids (Quantifoil Micro Tools), blotting for 4.5 seconds, and then plunge-freezing in liquid ethane using a Mark IV FEI Vitrobot (Thermo Fisher Scientific). The grids were screened on a 200 keV Glacios electron microscope (Thermo Fisher Scientific) equipped with a K3 direct detection camera (Gatan) at magnification of 45,000x, physical pixel size of 0.86 Å, and total dose of 50 e-/Å². Final data were collected on a 300 keV Titan Krios (Thermo Fisher Scientific) with a K3 direct detection camera (Gatan) and an energy filter (Gatan) either at the Yale CryoEM Resource (YCR) facility with magnification of 81,000x, physical pixel size of 1.068 Å, energy filter slit width of 20 eV, and total dose of 50 e-/Å² or the Brookhaven National Laboratory Laboratory for BioMolecular Structure (LBMS) facility, with magnification of 81,000x, physical pixel size of 1.07 Å, energy filter slit width of 20 eV, and total dose of 50 e-/Å². Tomography data were collected with magnification of 64,000x, physical pixel size of 1.346 Å, on the Titan Krios at YCR facility with dose symmetrical scheme +/− 54°, 3° increment and grouped by 3 on a starting tilt angle of 0° using PACE-tomo scripts^44^.

### Data processing and model building

Cryo-EM data processing was performed using CryoSPARC v4.4.1^45^. Raw movies were motion-corrected and CTF-corrected using CryoSPARC’s implementation. Initial particles were manually picked, followed by 2D classification to generate templates for template-based particle picking ^45^. Side view and top view classes were used in separate template-based particle picking jobs for efficient picking of both views. For datasets with pixel size of 1.068 Å collected on the Krios microscope, initial particles were extracted with a box size of 282 pixels and Fourier cropped to 64 pixels, while for data sets with pixel size of 0.86 Å, collected on a Glacios microscope, a box size of 346 was used and cropped to 128 pixels. Iterative cycles of heterogeneous refinement were performed with reference volumes, including a previously resolved map of the immature CA lattice (EMD-45975)^27^ and three copies of a featureless flat-ellipsoid-shaped map to absorb noise particles into junk classes. C6 symmetry was applied in heterogeneous refinements. Upon convergence after several rounds of heterogeneous refinement, the well-defined class of particles was re-extracted with a box size of 282 pixels (or 346 for 0.86 Å pixel size) without cropping. Re-extracted particles were then refined with local refinement. 3D classification and subsequent local refinements were performed to select the final well-defined particle set. This final particle set underwent global and local CTF refinements and reference-based motion correction, followed by a final round of local refinement^46^. The local resolution range was analyzed using a local resolution job (**Table S1 and Table S2**). Detailed processing steps for each dataset are illustrated in supplemental figures (**Fig. S7, S8, S9, S10**).

For *in situ* VLP data processing, we used the 2D template matching implemented in GisSPA^47^, which is particularly effective for low signal-to-noise *in situ* datasets^48, 49^. To rule out template bias, we used a “reference-omission” strategy: for CA6 (CA-NC), we omitted density corresponding to all six copies of the CA-SP1 six-helix bundle (CA residues 223-234 and all SP1 residues), and for CA5 (CA-SP1) we omitted the same region plus density corresponding to all six copies of residues lining the CA_NTD_ pore (residues 1–28). GisSPA-picked particles were then imported into cryoSPARC for 3D classification, subset selection using per-particle scale factors, and iterative local refinement with global and local CTF refinement. Only particles from classes that faithfully recovered the omitted regions were retained as true particles, whereas false positives produced reconstructions that maintain the omission pattern of the template. This workflow enabled robust identification of the rare hybrid lattice directly from intact *in situ* VLPs, including CA-SP1 and CA-NC samples (**Fig. S8, S9**), and supported efficient processing of IP6-treated perforated VLPs (**Fig. S10**).

Model building was performed by docking the initial model in ChimeraX^50^, followed by rebuilding and refinement using Coot^51^ and Phenix real space refinement^52^. Representation of the model with B factor coloring was rendered in ChimeraX. The average B factor of each residue or motif was calculated in the baverage program in the CCP4 program suite^53, 54^.

### Cryo-ET data processing and subtomogram averaging

Cryo-ET data processing and subtomogram averaging were primarily carried out following the RELION 5 workflows^55^. Cryo-EM movies were motion-corrected using MotionCor2^56^ and CTF estimation with CTFFIND4^57^. For denoising, tilt series were separated into odd and even tilts during motion correction, and the resulting stacks were processed using Cryo-CARE^58^. Tilt series were aligned with AreTomo2^59^, and CTF-corrected tomograms for particle picking were reconstructed with RELION5.0 at a pixel size of 12 Å. A total of 181 tilt-series were selected for further processing. VLPs are annotated on denoised tomograms using Napari^60^ implemented in RELION5 in sphere mode, and initial particle positions were uniformly distributed over the defined spherical surfaces with a spacing of 60 Å, resulting in 453,317 starting positions. For each extraction position, the Euler angles were defined to orient the extracted volumes normal to the VLP surface.

Particles were extracted in RELION5 and saved as 2D stacks. Three rounds of 3D classification were then performed to separate different capsid conformations. A total of 50,093 particles were identified as immature capsid, and 31,532 particles were retained after removing duplicates. An additional three rounds of 3D classification were conducted on 13,171 particles to isolate the hybrid capsid conformation, yielding 2,392 particles with an estimated resolution of 10.6 Å. For the immature conformation, particles were refined in one round in RELION5 to a resolution of 6.4 Å. The resulting particle sets were then mapped back onto the tomograms in Napari to analyze their 3D distribution on the VLP surfaces. Detailed processing steps for the dataset are illustrated in a supplemental figure (**Fig. S11**).

### Virus production and quantification

HIV-1 stocks were produced by polyethyleneimine-mediated transfection of 293T cell monolayers with the wild type full-length HIV-1 molecular clone R9^61^ and the mutant encoding the H84A substitution in CA. After overnight culture, the supernatant was removed and the cells rinsed with 5 mL PBS, then replenished with 6 mL of complete medium (DMEM supplemented with fetal bovine serum (10% vol/vol) and penicillin plus streptomycin. Culture supernatants were harvested 24-32 hours later. A one mL volume of each virus stock were treated with DNAse I (20 mg/mL) in the presence of 10 mM MgCl_2_ for one hour at 37°C. Virions were then pelleted by ultracentrifugation at 4°C through a 0.25 mL cushion of STE buffer (10 mM Tris-HCl pH 7.5, 150 mM NaCl, 1 mM EDTA) containing 20% sucrose (wt/vol) using a Beckman TLA-55 rotor. The supernatant was carefully removed by aspiration, and the pellet was resuspended in 50 µL STE buffer by gentle but thorough pipetting and the solution transferred to a clean microfuge tube. Lysates of the transfected cells were prepared by rinsing the monolayers with 5 mL PBS, detachment by scraping into 1 mL of PBS, pelleting, and resuspending in 200 µL of STE buffer containing 1% Triton X-100. Virus stocks were quantified by assaying exogenous RT activity using a previously described radiometric assay^62^.

### ERT assays

ERT and NERT assays were performed as described^63^. Briefly, 5 μl of concentrated, DNAse I-treated virus suspension was added to a reaction mix containing 20 mM Tris-HCl pH 7.6, 150 mM NaCl, 2 mM MgCl_2_, 1 mg/mL BSA, 0.1 mM each dNTP, and 0.5 mM DTT containing (ERT reaction) or lacking (NERT reaction) Triton X-100 (0.1% vol/vol) and inositol hexakisphosphate (IP6, 100 mM). Following incubation at 37°C for 16h, DNA products were purified on silica spin columns (Epoch Life Science), eluted with 100 μl of water, and assayed for early (minus strand strong stop) and late (full length minus) products by qPCR using Taqman chemistry in a Stratagene 3000p real time thermal cycler. Values were interpolated from standard curves generated from serial dilutions of R9 plasmid DNA.

### Assay of HIV-1 infectivity

Concentrated virus stocks were diluted in complete medium and inoculated onto monolayers of TZM-bl cells in a 96 well plate. After 2 days of culture, luciferase activity in cell lysates was quantified in a Molecular Devices L-max 96-well luminometer following addition of 30 μL of lysis buffer containing 50 mM Tris-HCl (pH 7.8), 130 mM NaCl, 10 mM KCl, 5 mM MgCl_2_, and 0.5% Triton X-100. Following lysis, each well was injected first with 200 μL of a solution of 75 mM Tris HCl (pH 8.0) containing 8.3 mM magnesium acetate and 4 mM ATP and subsequently with 80 μL of a solution containing 1 mM D-luciferin potassium salt (Gold Biotechnology, cat.no. eLUCK-1G). Luminscence signals were integrated for 5 seconds. Infectivity was determined as the relative light unit (RLU) value normalized by exogenous RT activity in the respective virus inoculum.

### Molecular Dynamics Simulations Preparation

Atomistic models for the immature (PDB:9CWV)^27^, hybrid (this work), and mature (PDB:8CKV)^39^ CA lattices were derived from experimental structural data. Subunit interface transitions were examined using selections representing the hexamer, trimer, and monomer. The N-terminal (P1-H12) and C-terminal (T243-M245) tails were modeled with Modeller (v.10.4) implemented in ChimeraX^50, 64^. Residues from the HXB2 sequence in the immature and hybrid structures were remodeled to match the pNL4-3 (WT) sequence (L83V, H120N, G208A, P241S), and the T8I mutation in SP1 was reverted to the WT sequence. Each system was solvated with TIP3 water molecules^65^ using orthorhombic boxes generated with the VMD solvate plugin^66^, ionized to 150 mM NaCl with charge neutrality via the VMD autoionize plugin^66^ (**Table S3**). Care was taken such that the water boxes were made large enough for any domain transitions or movement would not come near the edges. Simulations were visualized and analyzed using VMD and in-house TCL scripts^67^.

For targeted molecular dynamics (TMD) preparation, solvent was first minimized for 20,000 steps via the gradient descendent algorithm with the protein backbone fixed. To ensure the CA NTD-CTD linker was sufficiently relaxed, an iterative minimization procedure (20 iterations) was carried out: (1) the CA_NTD_ and CA_CTD_ backbones were restrained (10 kcal/mol Å) with the linker free, and (2) the protein domains were free while the linker backbone was restrained (10 kcal/mol Å). The solvent was then gradually heated to 310K in 5K increments with the protein backbone fixed, followed by a final step where the backbone was harmonically constrained (10 kcal/mol Å) and a final thermalization step following the mentioned procedure with the same backbone constraint. The harmonic constraints were then gradually removed in an NPT simulation at 310K and 1 bar. Targeted molecular dynamics were then run for each assembly transition, as described below, with harmonic constraints (10 kcal/mol Å) applied to the helical Cα in CA_CTD_ to maintain an appropriate domain reference and appropriate lattice assembly effects.

All simulations were carried out using NAMD3^68^ with CHARMM36m forcefield parameters for protein, water, and ions^69–72^. The temperature was controlled with the Langevin thermostat with a damping coefficient of 0.1ps^-1^ and the pressure was controlled with the Nose-Hoover Langevin piston method to 1 bar with periodic boundary conditions. Hydrogen bonds were constrained with SHAKE^73^ and SETTLE^74^ algorithms. Non-bonded interactions were calculated with a cutoff distance of 12 Å for electrostatic interactions with the Particle-Mesh-Ewald^75^ summation with a grid size of 1.0 Å.

### Targeted Molecular Dynamics

To drive the transition from the starting structure to the target structure, we employ appropriate colvar functions for TMD of each transition^76, 77^. Standard TMD employs an RMSD-based algorithm to drive the coordinates along an arbitrary linear interpolation pathway between two structures. The hybrid to mature transition is dominated by a translational movement of the CA_CTD_, corresponding to a ∼40 Å expansion, to which the RMSD-based algorithm is sufficient to describe the motion. To preserve CA_NTD_-CA_NTD_ interfaces during the transition, harmonic constraint (1.0 kcal/mol Å) were applied to the Cα of the CA_NTD_ helices. The RMSD between the CA_CTD_ of the hybrid and mature monomers were measured at 27.6 Å when the CA_CTD_ regions are superimposed. To optimize the biasing strength, we performed a scan of force constants over 10 ns trajectories and determined that 2.0 kcal/mol Å was the minimal constant sufficient to drive the complete transition, which was then used for further analysis.

In contrast, the immature to hybrid transition is dominated by a large CA_NTD_ rotation, making a linear RMSD interpolation unsuitable. Using the angle between the Cα atoms of E29-P147-Q176 as a reference, the immature protomer measures ∼47^◦^, whereas the hybrid protomer measures ∼230^◦^, corresponding to a rotation of nearly 180^◦^ (**Fig. S5A**). A direct linear interpolation of such a rotation distorts the tertiary structure. Instead, we used the orientation colvar function to describe the CA_NTD_ rotation^78, 79^. A limitation of quaternion-based colvars is that the rotation always follows the shortest arc pathway between the two orientations. For the immature to hybrid transition, the shortest arc corresponds to ∼140^◦^ in the clockwise direction (**Fig. S5B**). Moreover, quaternions become ambiguous as it approaches 180^◦^, so to probe the counterclockwise pathway we instead reversed the process, driving the transition from the hybrid to the immature state.

To preserve CA_CTD_-CA_CTD_ interfaces during the immature to hybrid transition and establish a stable reference frame, we applied a harmonic constraint (1.0 kcal/mol Å) to the Cα of the CA_CTD_ helices. We then performed a sweep of constant values (3,286 kcal/mol×rad^2^ = 1 kcal/mol×deg^2^) on the orientation colvar to identify the minimal bias required to complete the domain rotation within 10 ns. Completion was defined by reasonable alignment of the domain with the target structure, monitored via RMSD. For clockwise rotation, a force constant of 1.0 kcal/mol×deg^2^ produced a rapid transition, whereas lower constants left the CA_NTD_ trapped near its starting orientation. For the counterclockwise pathway, the first half of the rotation occurred with constants as low as 0.02 kcal/mol×deg^2^; however, the full transition could not be completed, even up to constants of 3.0 kcal/mol×deg^2^. We therefore selected 0.9 kcal/mol×deg^2^ for further analysis, despite the incomplete rotation.

We then tried to rotate the CA_NTD_ of the C6 subunit which contains a full hexamer as well as the C3 subunit which is a trimer of the interface between three hexamers, using the same CA_CTD_ constraints and the same target quaternion for each transition. In a similar scan, we apply the force simultaneously to all six CA_NTD_ in the hexamer as well as one CA_NTD_ for the C6 subunit. In the simultaneous transition at a force constant of 0.4 kcal/mol×deg^2^, all six protomers are not observed to engage; however, one protomer is observed to continue with the rotation. We note the force constant used to capture the rotation of one protomer in the C6 rotation is lower than the one required to rotate the CA_NTD_ in the clockwise pathway. It is not observed to complete the full transition of all six CA_NTD_ within the allotted 10 ns. The protomeric transition is observed at 0.4 kcal/mol×deg^2^. For the C3 subunit, when a force is applied to all three or two CA_NTD_ simultaneously, the result is that they cannot complete the transition without deforming one or more CA_NTD_. Strikingly, when only applied to one of the three CA_NTD_ at a time, the CA_NTD_ is observed to complete the rotation we observed in the monomer in a clockwise path at a force constant of 0.4 kcal/mol×deg^2^.

## Acknowledgements

This work was funded in part by the National Institutes of Health through grants R01AI192025 (Y.X. and C.A.), U54AI170791 (Y.X., C.A., and J.R.P.), R01AI178846 (Y.X., C.A., and J.R.P.), and 1F31AI181652-02 (C.F.). Cryo-EM data was collected with the support of staff at the Brookhaven National Laboratory – Laboratory for BioMolecular Structure (LBMS) and the Yale CryoEM Resource (YCR). LBMS is supported by the DOE Office of Biological and Environmental Research (KP1607011). Yale YCR is funded in part by the NIH grant S10OD023603. We thank the Xiong lab members for discussions. We acknowledge the computational resources provided by the Delaware Advanced Research Workforce and Innovation Network (DARWIN) as well as the Caviness cluster. This work used Stampede3 at TACC, Delta at NCSA, and Anvil at RCAC Purdue through allocation MCB-170096 from the Advanced Cyberinfrastructure Coordination Ecosystem: Services & Support (ACCESS) program, which is supported by National Science Foundation awards (2138259, 2138286, 2138307, 2137603, and 2138296).

## Author contributions

M.E.M, C.W., and Y.X. conceived the concept and drafted the initial manuscript. M.E.M., R.R.Y., J.S. and C.A. prepared the viral-like particle samples and performed assays of HIV-1 infectivity, and ERT and NERT activity. C.F. prepared the liposomes. M.E.M and R.R.Y. prepared the cryo-EM samples, and together with C.W. performed cryo-EM grid screening. C.W. and J.L. collected the cryo-EM and cryo-ET data. M.E.M, C.W., and Y.X. performed cryo-EM reconstruction and analysis. J.C. carried out the tomography analysis. J.R.P. and A.M. designed and performed the molecular simulation and interpreted the results. M.E.M., C.W., C.A., J.R.P. and Y.X. took lead in writing the manuscript and all authors discussed and contributed to the manuscript preparation.

## Materials and Correspondence

Correspondence and material requests should be addressed to Y.X.

## Data Availability

Maps generated from the electron microscope data are deposited in the Electron Microscopy Data Bank under EMD-XXX. Atomic models have been deposited in the RCSB PDB with PDB IDs XXXX.

## Supplemental materials

**Figure S1.**
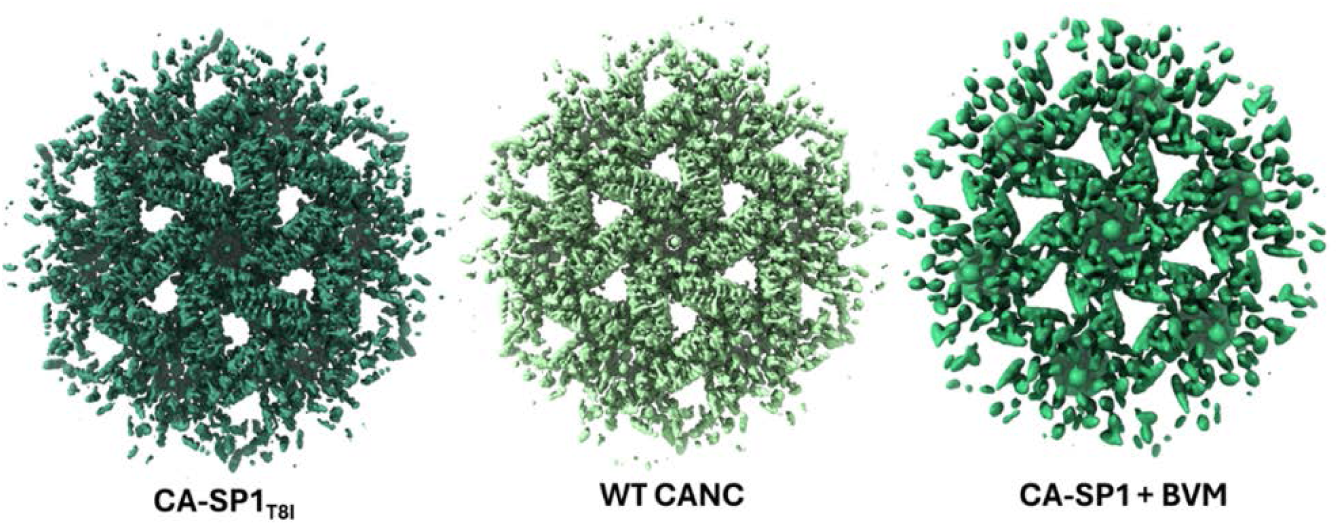
Cryo-EM maps of *in vitro* hybrid lattice assemblies. Samples include large unilamellar vesicle (LUV)-templated CA-SP1_T8I_ (forest green) (2.58 Å), WT CANC (sea green) (2.87 Å), and LUV-templated CA-SP1 treated with BVM (spring green) (3.74 Å).

**Figure S2.**
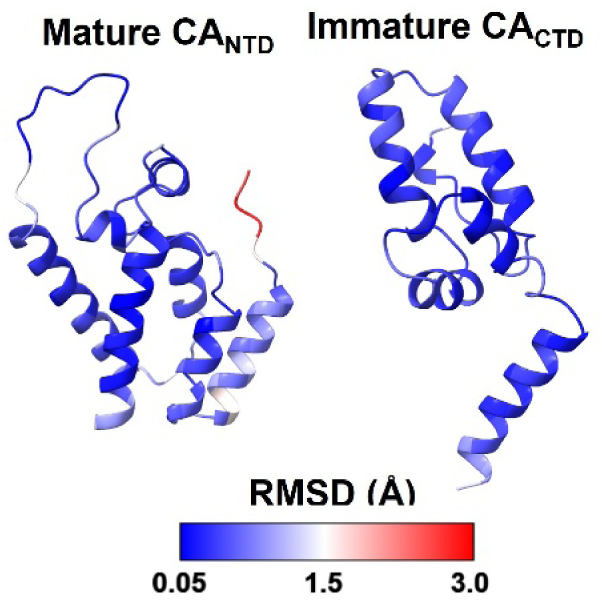
RMSD analysis of the hybrid CA_NTD_ and CA_CTD_-SP1 relative to corresponding immature and mature conformations. Comparison of the mature CA_NTD_ to the CA-SP1_T8I_ hybrid CA_NTD_ (left) and the immature CA_CTD_ to the CA-SP1_T8I_ hybrid CA_CTD_ (right), illustrating the structural similarities within these domains.

**Figure S3.**
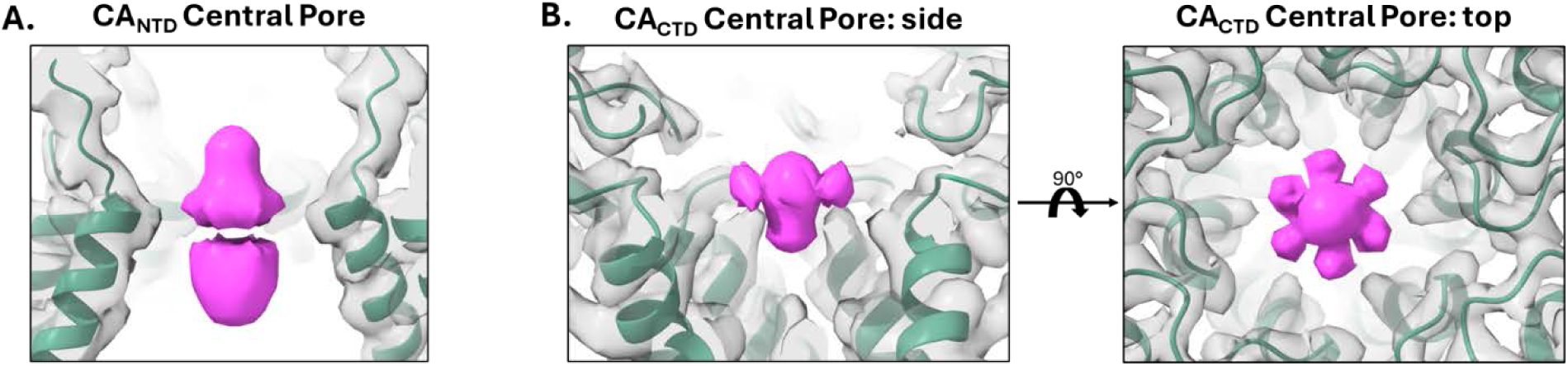
The hybrid lattice binds three IP6 molecules. A. Two IP6 molecules (magenta) bind to the CA_NTD_ of the hybrid lattice (green). The upper one is coordinated by R18 and the lower one is coordinated by K25. B. One IP6 molecule (magenta) binds to the CA_CTD_ of the hybrid lattice (green). The IP6 molecule is coordinated by K159 and K227 and is shown from side (left) and top (right) views.

**Figure S4.**
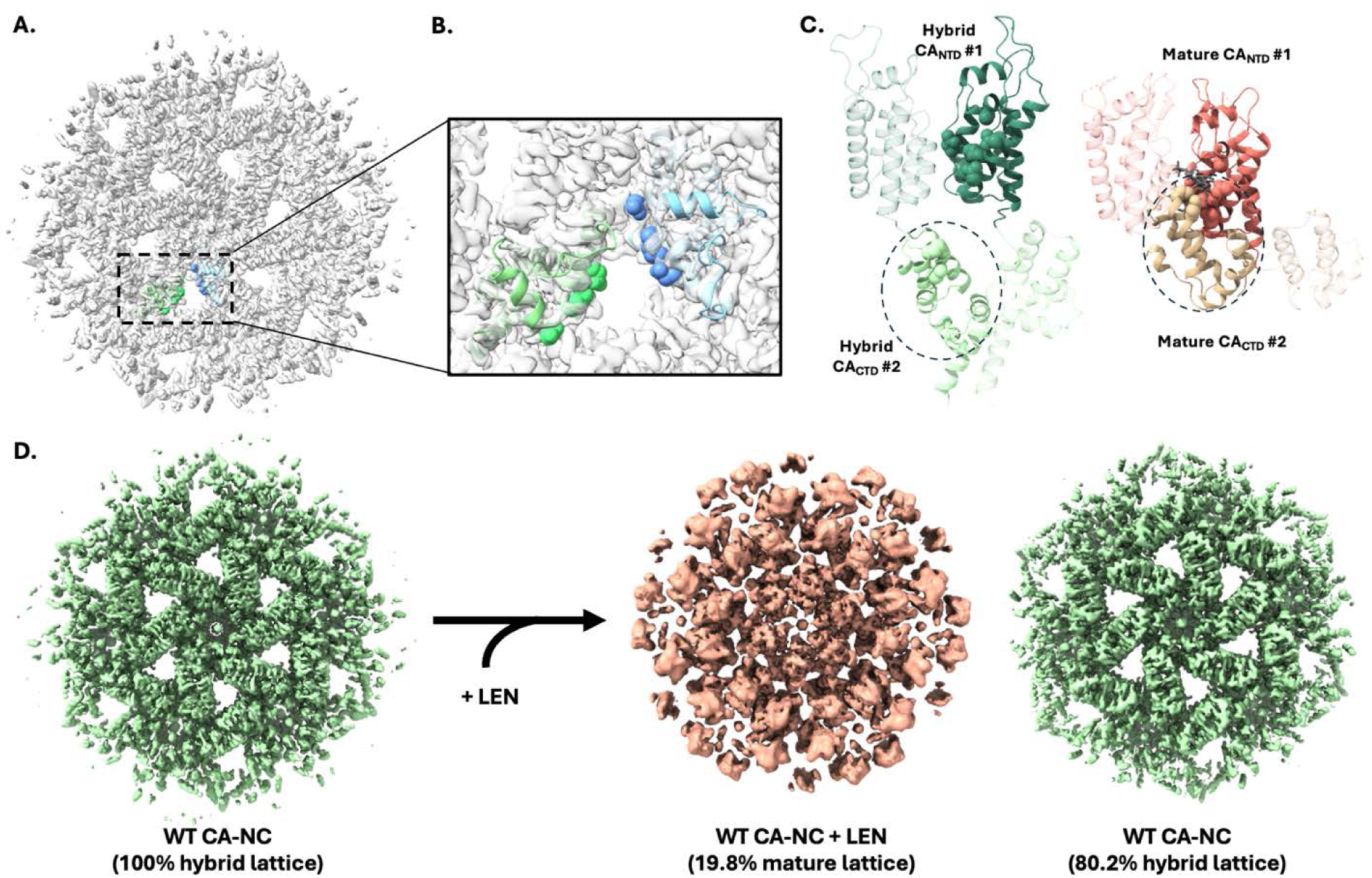
LEN binding shifts hybrid lattice into mature lattice. A. LEN-binding residues on chains A (blue) and B’ (mint) are exposed to the solvent accessible channel in the hybrid lattice. B. Residues N74, K70, and N57 are shown in spheres on chain A (blue) and chain B’ (mint). C. Comparison of FG-binding pocket organization in hybrid and mature lattices. Left, in the hybrid lattice, the FG-binding pocket lacks stabilizing “capping” interactions from the neighboring CA_CTD_ (#2) due to its extended conformation, leaving the pocket incompletely formed. Right, in the mature lattice, the FG-binding pocket is capped by the neighboring CA_CTD_ (#2), forming a stable binding site. LEN-binding residues are shown as spheres. CA_NTD_ is colored dark green (hybrid) or salmon (mature), and CA_CTD_ is colored light green (hybrid) or light pink (mature). CA chains and domains that are not involved in forming the highlighted FG-binding pocket are transparent. LEN is shown in the mature FG-binding pocket in grey. D. Left, cryo-EM analysis of WT CA-NC (cryo-EM map in green) yields a population of 100% hybrid lattice. Right, upon treatment with LEN, the WT CA-NC population becomes a mixture of 19.8% mature lattice (pink, left) and 80.2% hybrid lattice (mint, right).

**Figure S5.**
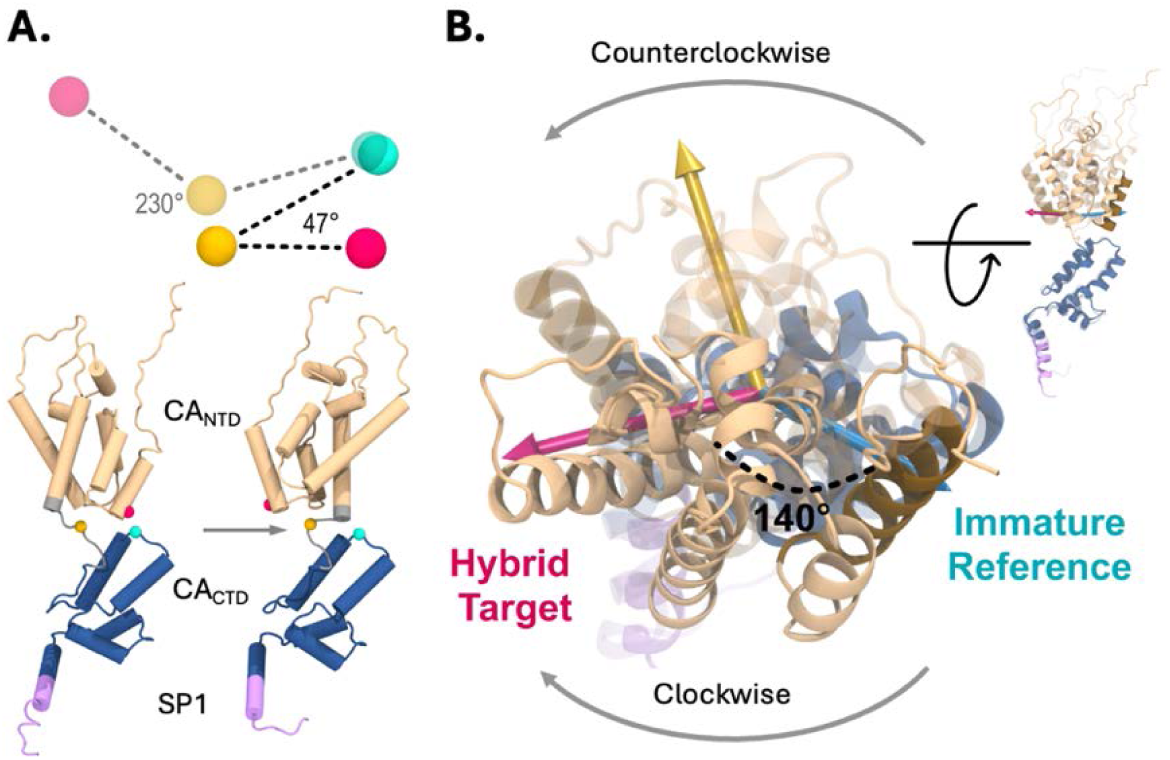
Diagram illustrating the measured (A) E29-P147-Q176 Cα angle (magenta-gold-cyan) and (B) quaternions used to transverse the CA_NTD_ rotation during the transition between the immature and hybrid states. The hybrid monomer is shown in a transparent view to depict the target of the rotation. The identity quaternion of the immature monomer is shown in light blue, while the target quaternion of the hybrid is shown in pink. The short arc along the quaternion-defined path is 140^◦^ rotation in the clockwise direction. The complementary counterclockwise rotation is then characterized as 220^◦^.

**Figure S6.**
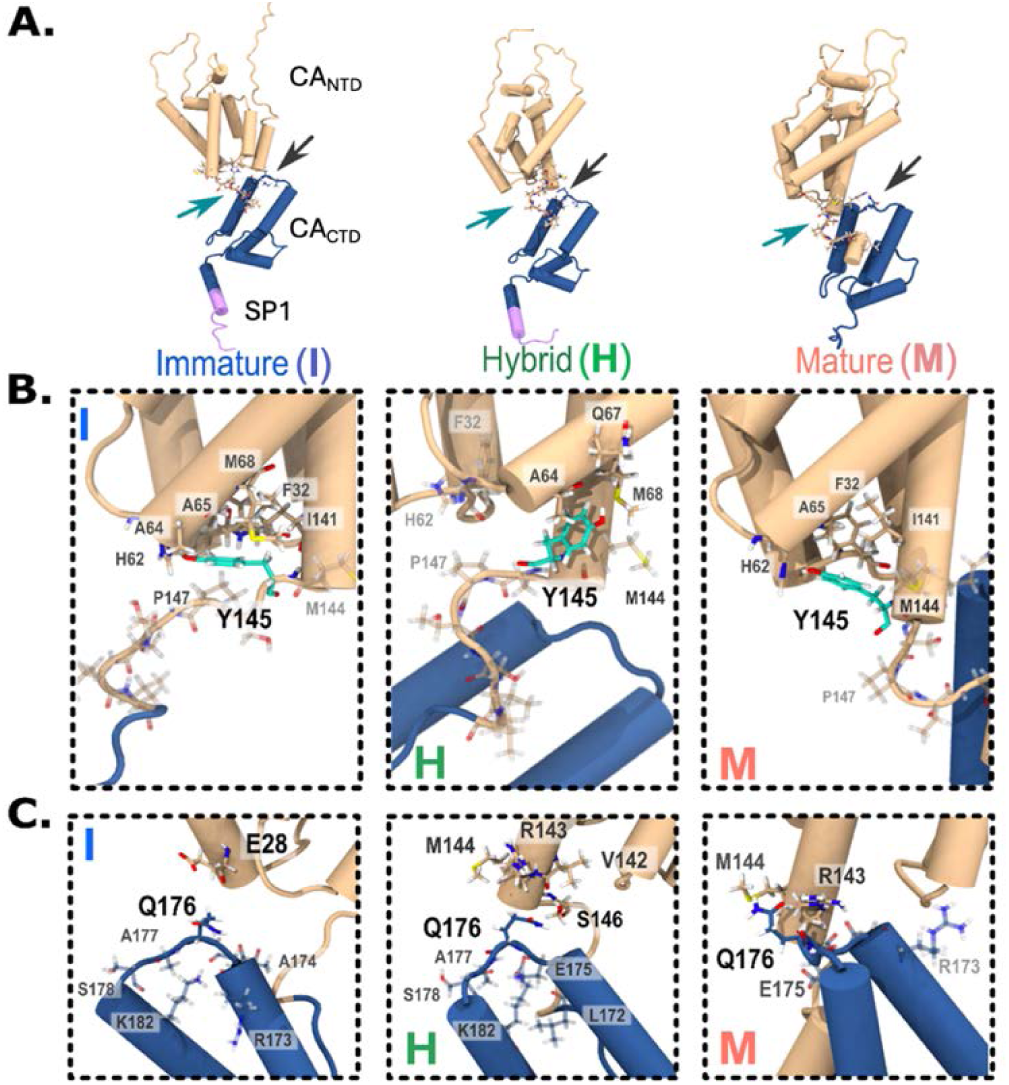
(A) Snapshots of the immature, hybrid, & mature CA highlighting the two regions of interest from the simulations: (B) changes in the CA NTD-CTD linker (around Y145), marked by cyan arrow in (A), and (C) CA_NTD_-CA_CTD_ (Q176) contacts, marked by black arrow in (A). Residues within 5 Å of these identified key residues are highlighted as the packing of sidechains is rearranged during the transition.

**Figure S7.**
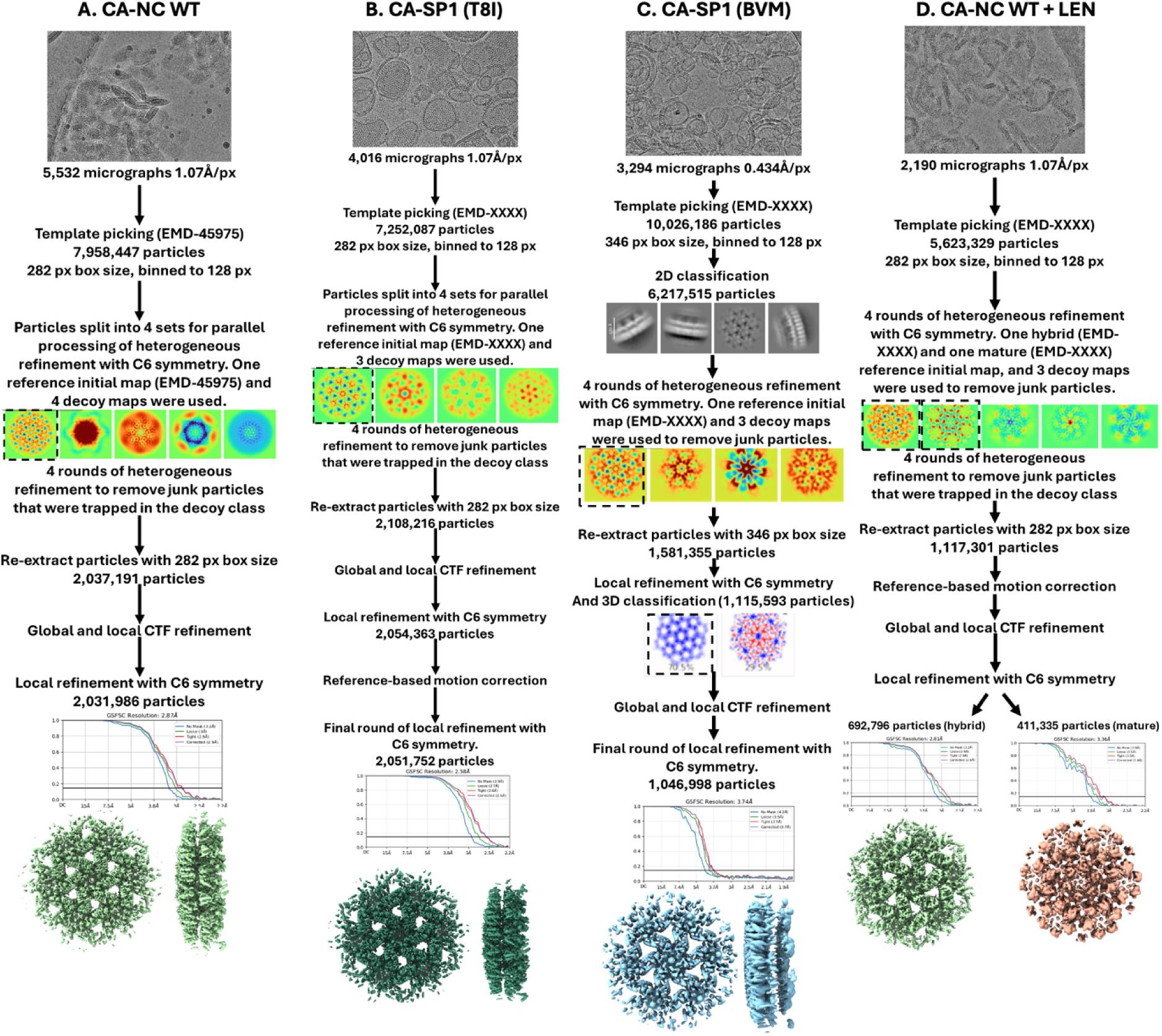
Data processing pipelines for *in vitro* assembled hybrid lattices. Each panel outlines the steps taken from raw cryo-EM micrographs through particle selection, classification, and 3D reconstruction to obtain the final cryo-EM maps for the respective lattice assemblies. A. Data processing workflow for CA-NC WT assembled with tRNA. B. Data processing workflow for CA-SP1_T8I_ assembled with LUVs. C. Data processing workflow for CA-SP1 treated with BVM and assembled with LUVs. D. Data processing workflow for CA-NC WT assembled with tRNA and treated with LEN.

**Figure S8.**
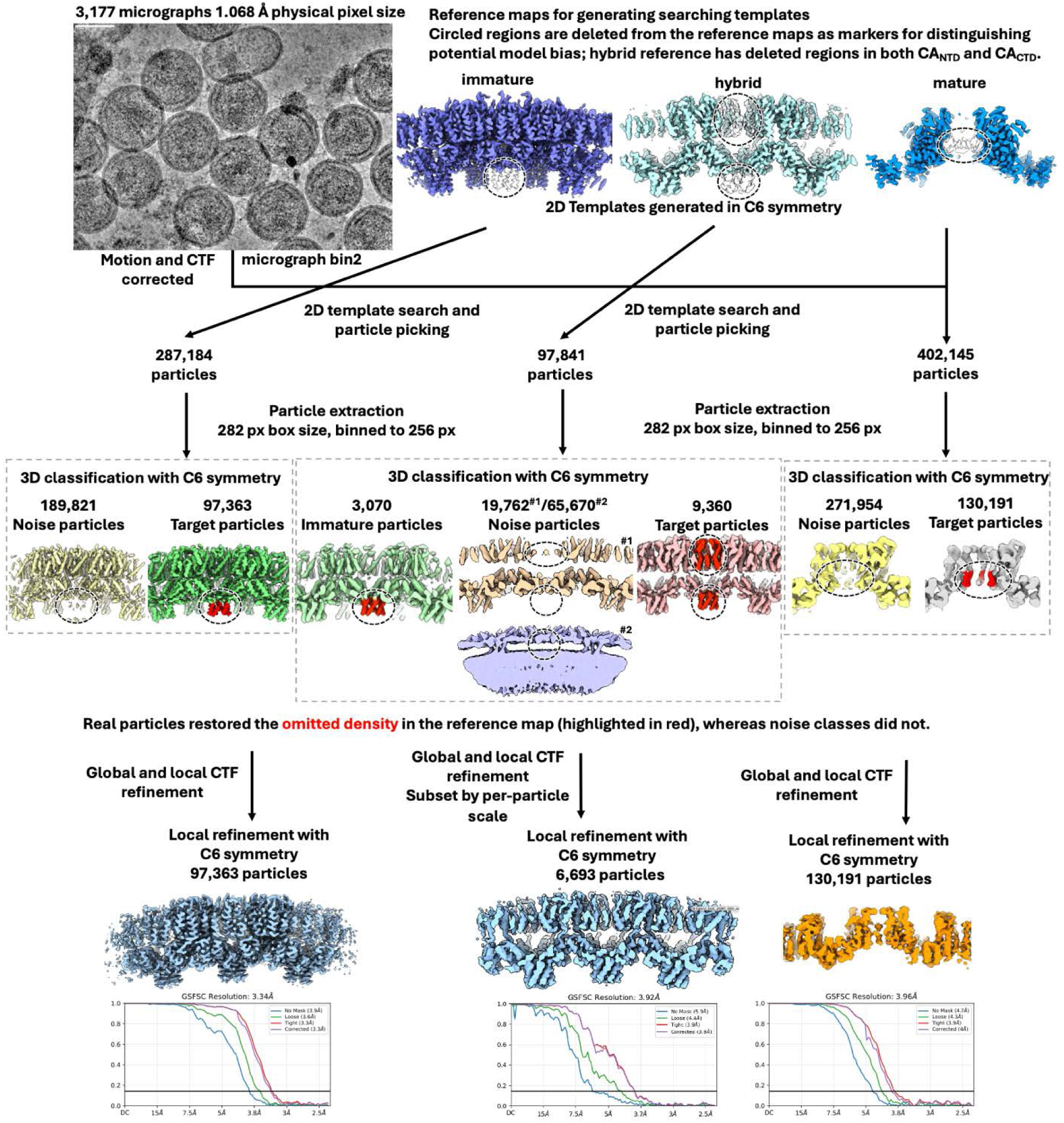
Data processing pipelines for *in situ* CA-SP1 VLP lattices. Data processing using 2D template matching using part-omitted (red) reference maps to reconstruct the immature (left), hybrid (center), and mature (right) lattices.

**Figure S9.**
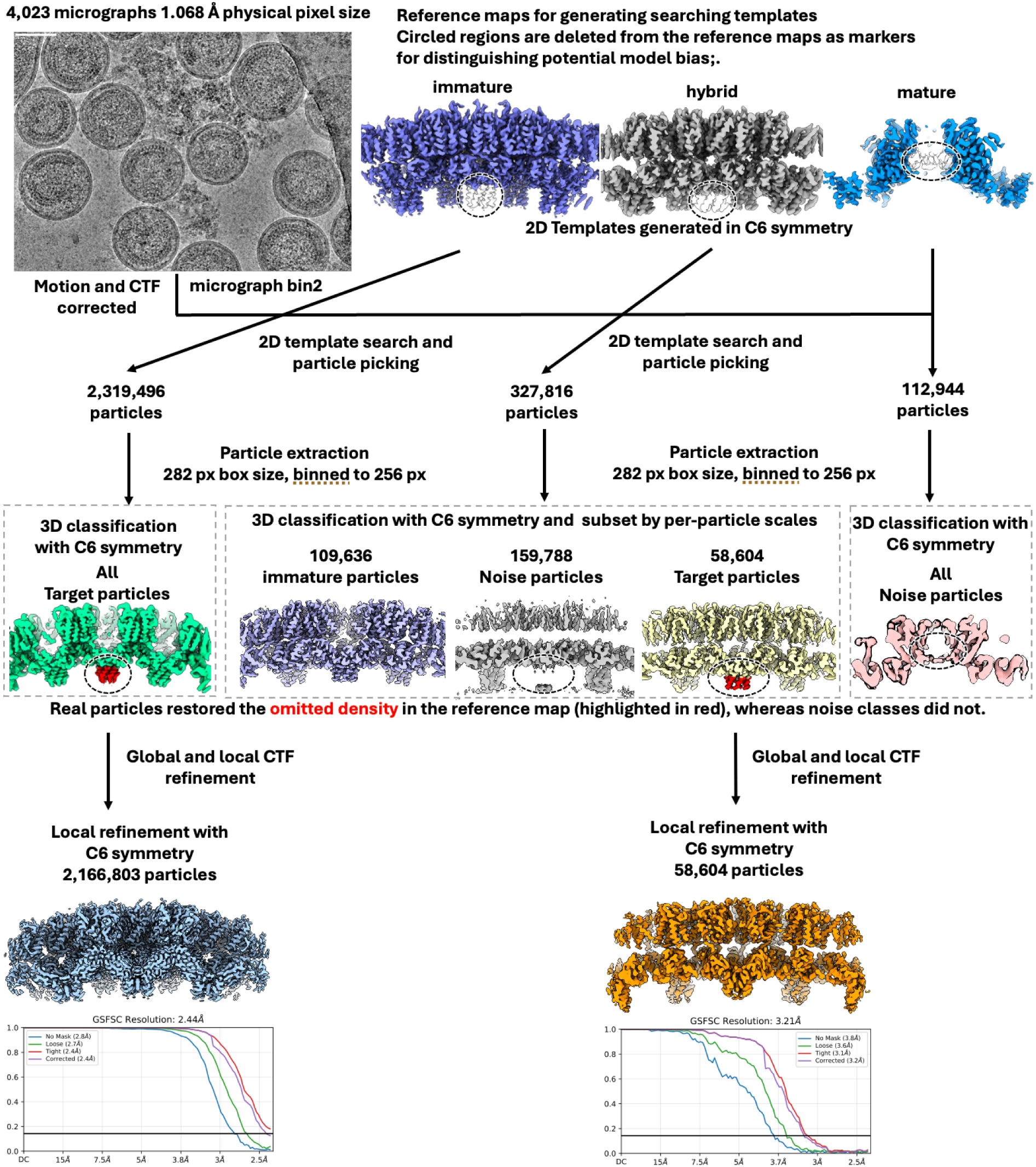
Data processing pipelines for immature and hybrid lattices from *in situ* CA-NC VLPs. Data processing using 2D template matching using part-omitted (red) reference maps to reconstruct the immature (left) and hybrid (right) lattices.

**Figure S10.**
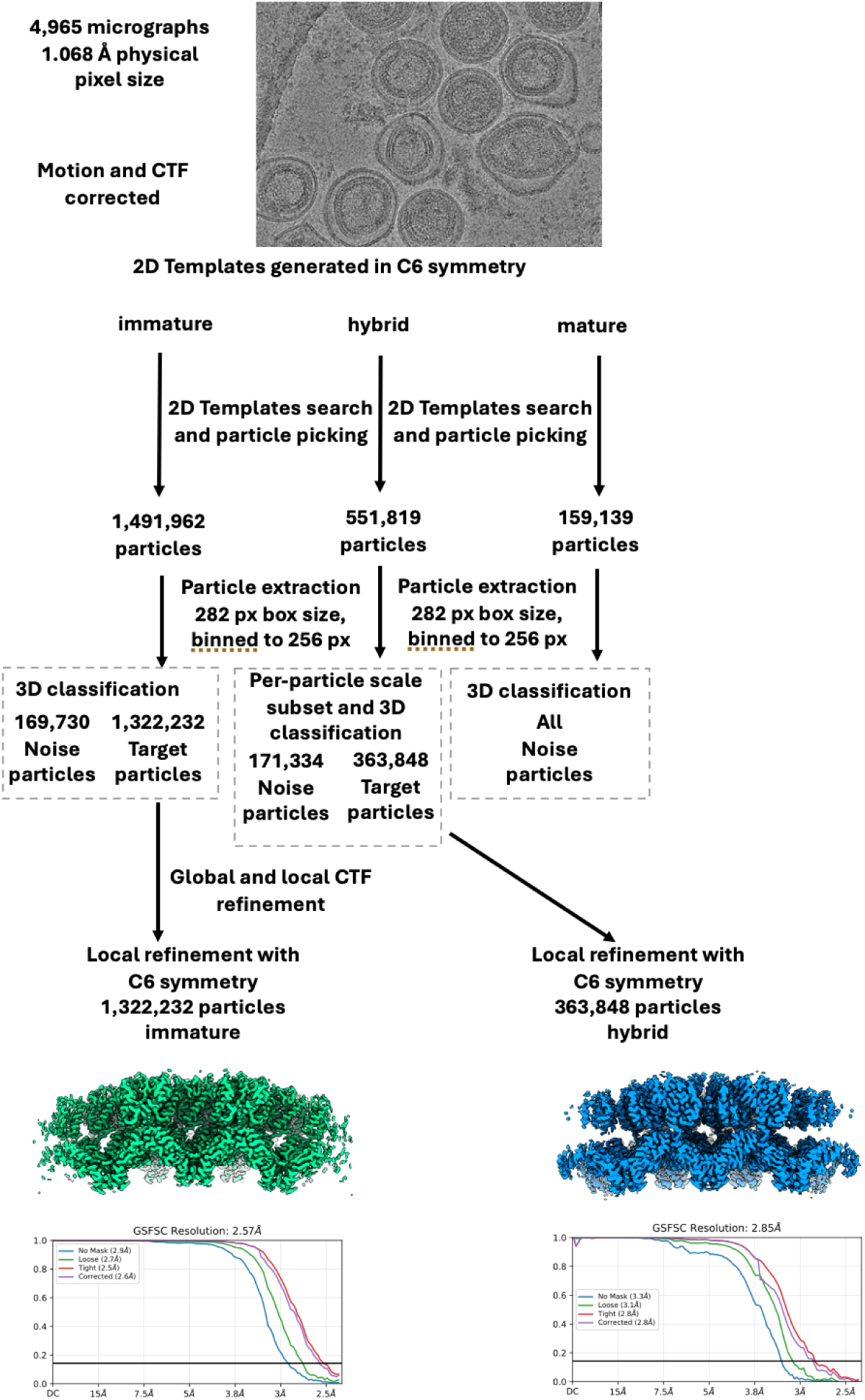
Data processing pipelines for immature and hybrid lattices from *in situ* CA-NC VLPs treated with IP6. Data processing using 2D template matching on CA-NC VLPs treated with PFO and IP6 to reconstruct the immature (left) and hybrid (right) lattices.

**Figure S11.**
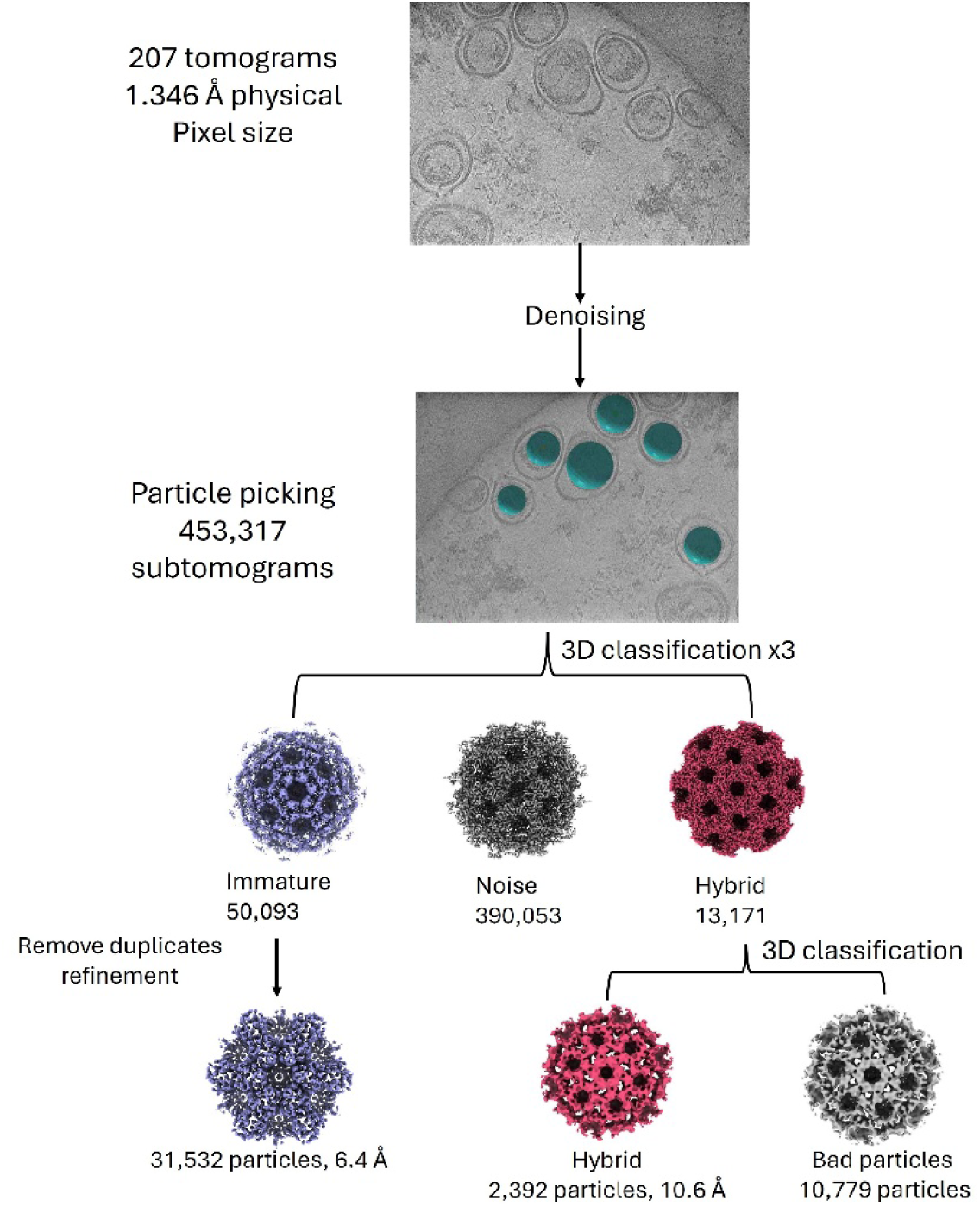
Cryo-ET data processing pipelines for immature and hybrid lattices from *in situ* CA-SP1_T8I_ VLPs treated with IP6. Data processing for CA-SP1_T8I_ VLPs treated with PFO and IP6 to reconstruct the immature (left) and hybrid (right) lattices.

**Table S1.**
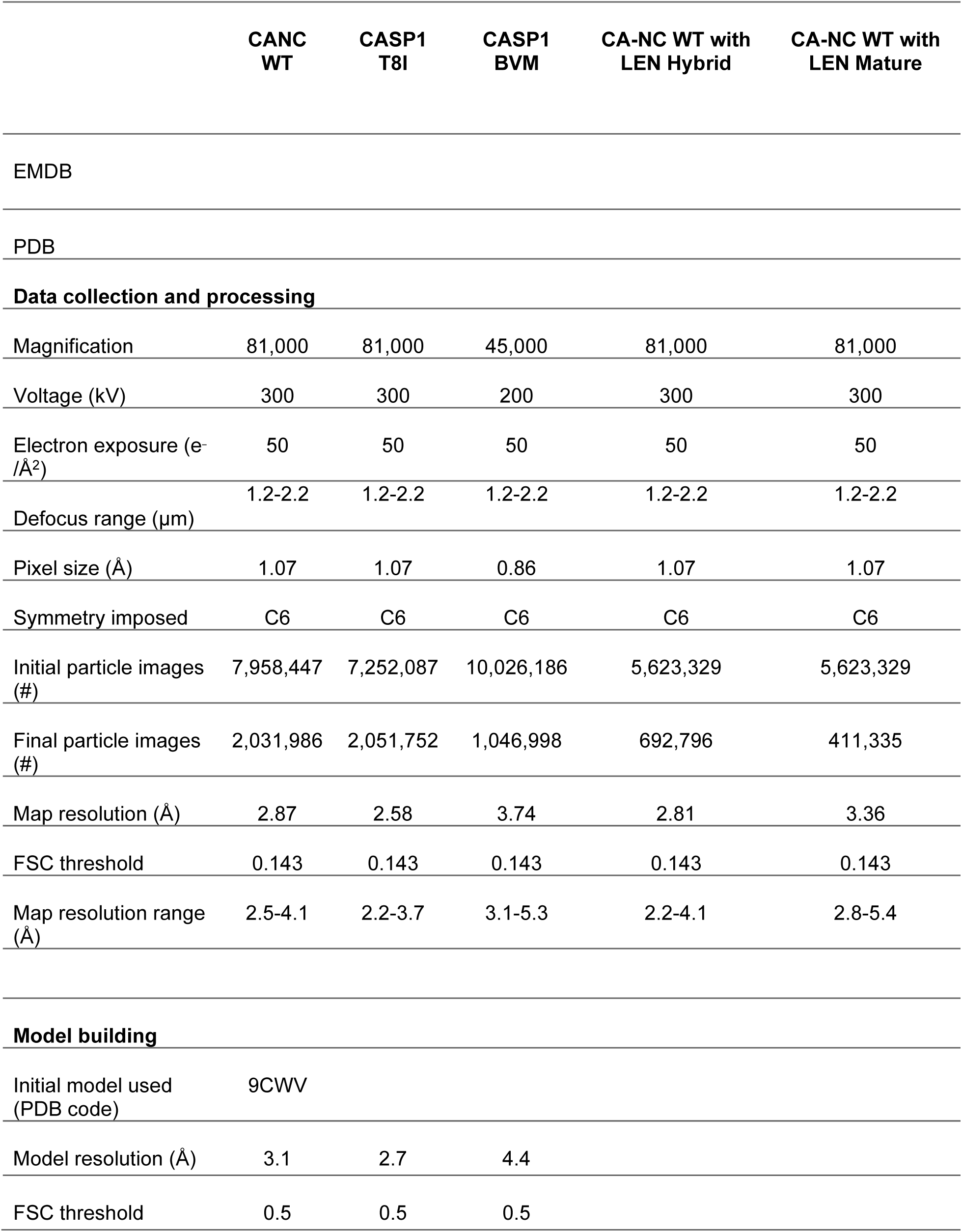
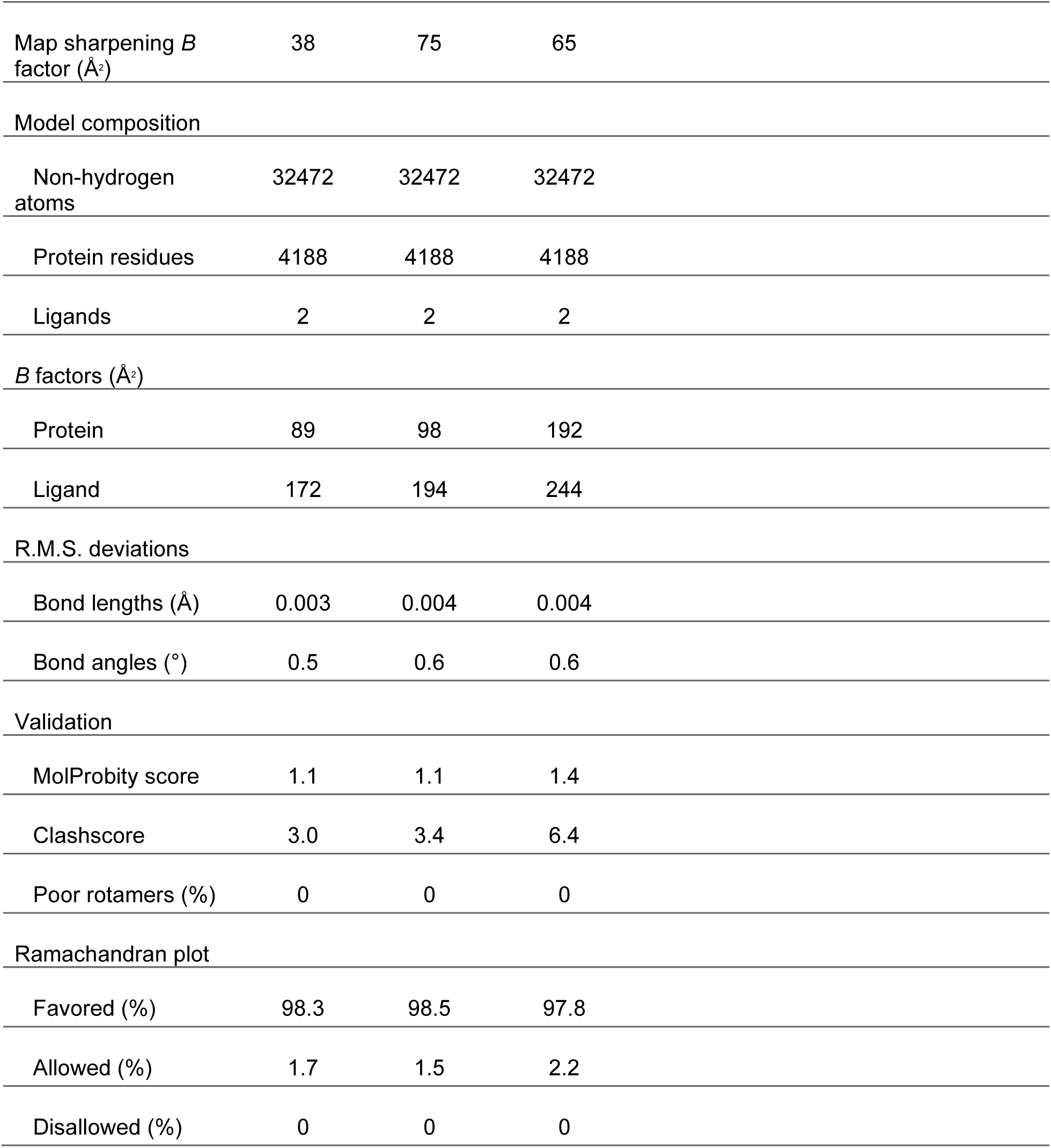
Cryo-EM data collection, refinement, and validation statistics for *in vitro* constructs.

|  | CANC<br>WT | CASP1<br>T8I | CASP1<br>BVM | CA-NC WT with<br>LEN Hybrid | CA-NC WT with<br>LEN Mature |
| --- | --- | --- | --- | --- | --- |
| EMDB |  |  |  |  |  |
| PDB |  |  |  |  |  |
| **Data collection and processing** |  |  |  |  |  |
| Magnification | 81,000 | 81,000 | 45,000 | 81,000 | 81,000 |
| Voltage (kV) | 300 | 300 | 200 | 300 | 300 |
| Electron exposure (e<sup>-</sup>/Å<sup>2</sup>) | 50 | 50 | 50 | 50 | 50 |
| Defocus range (μm) | 1.2-2.2 | 1.2-2.2 | 1.2-2.2 | 1.2-2.2 | 1.2-2.2 |
| Pixel size (Å) | 1.07 | 1.07 | 0.86 | 1.07 | 1.07 |
| Symmetry imposed | C6 | C6 | C6 | C6 | C6 |
| Initial particle images (#) | 7,958,447 | 7,252,087 | 10,026,186 | 5,623,329 | 5,623,329 |
| Final particle images (#) | 2,031,986 | 2,051,752 | 1,046,998 | 692,796 | 411,335 |
| Map resolution (Å) | 2.87 | 2.58 | 3.74 | 2.81 | 3.36 |
| FSC threshold | 0.143 | 0.143 | 0.143 | 0.143 | 0.143 |
| Map resolution range (Å) | 2.5-4.1 | 2.2-3.7 | 3.1-5.3 | 2.2-4.1 | 2.8-5.4 |
| **Model building** |  |  |  |  |  |
| Initial model used (PDB code) | 9CWV |  |  |  |  |
| Model resolution (Å) | 3.1 | 2.7 | 4.4 |  |  |
| FSC threshold | 0.5 | 0.5 | 0.5 |  |  |

|  |  |  |  |
| --- | --- | --- | --- |
| Map sharpening $B$ factor ($\text{\AA}^2$) | 38 | 75 | 65 |
| Model composition |  |  |  |
| Non-hydrogen atoms | 32472 | 32472 | 32472 |
| Protein residues | 4188 | 4188 | 4188 |
| Ligands | 2 | 2 | 2 |
| $B$ factors ($\text{\AA}^2$) | | | |
| Protein | 89 | 98 | 192 |
| Ligand | 172 | 194 | 244 |
| R.M.S. deviations |  |  |  |
| Bond lengths ($\text{\AA}$) | 0.003 | 0.004 | 0.004 |
| Bond angles ($^\circ$) | 0.5 | 0.6 | 0.6 |
| Validation |  |  |  |
| MolProbity score | 1.1 | 1.1 | 1.4 |
| Clashscore | 3.0 | 3.4 | 6.4 |
| Poor rotamers (%) | 0 | 0 | 0 |
| Ramachandran plot |  |  |  |
| Favored (%) | 98.3 | 98.5 | 97.8 |
| Allowed (%) | 1.7 | 1.5 | 2.2 |
| Disallowed (%) | 0 | 0 | 0 |

**Table S2.**
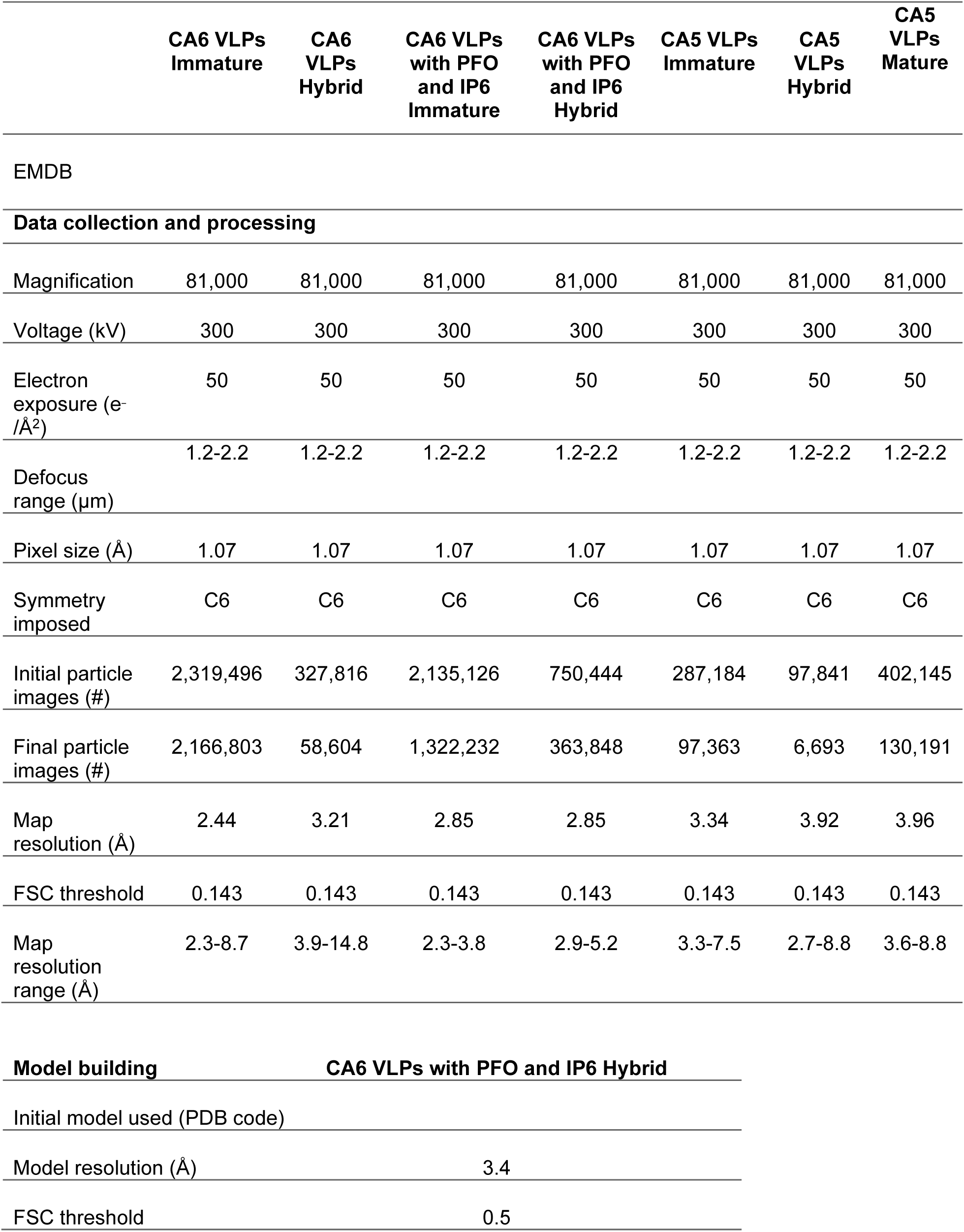
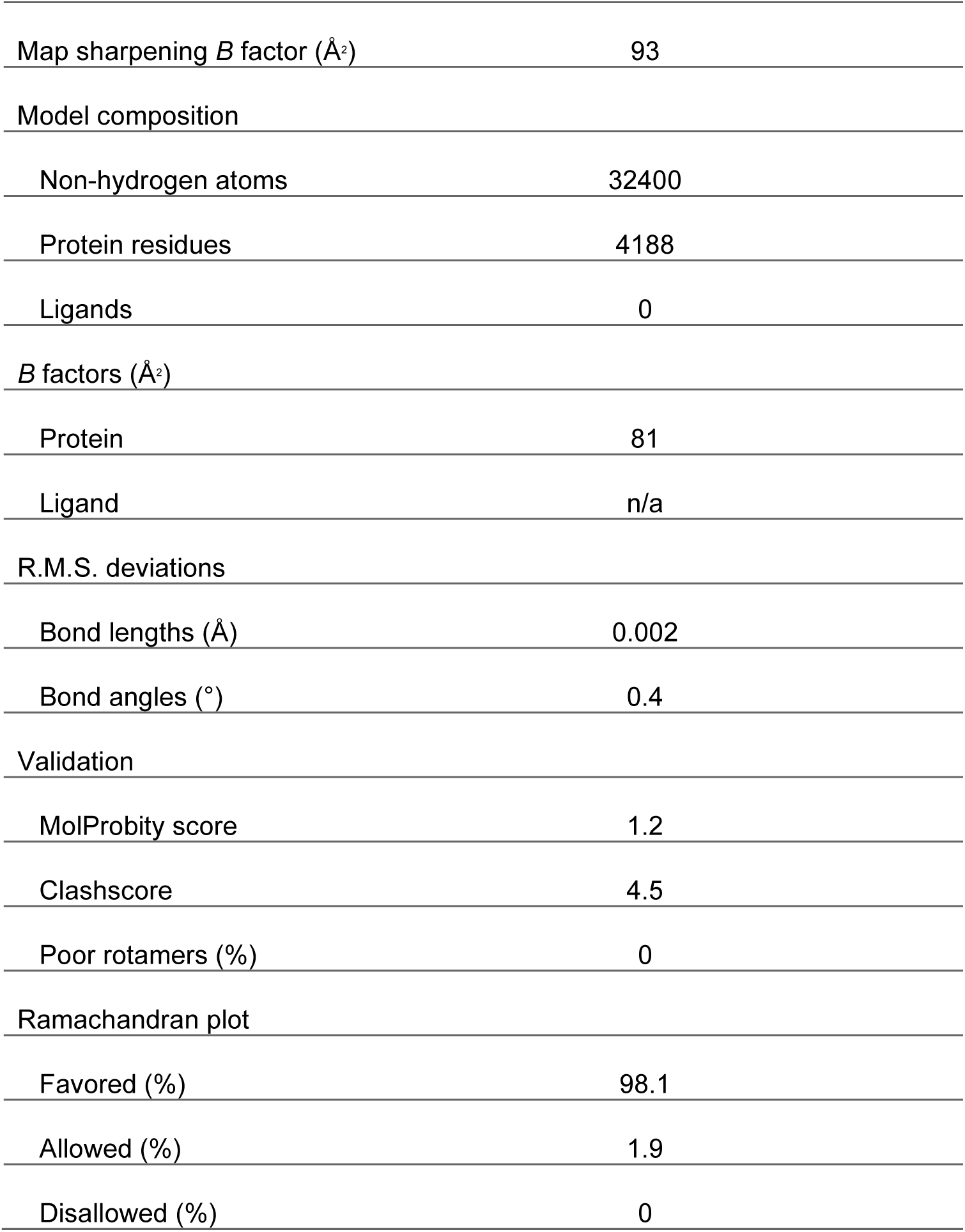
Cryo-EM data collection, refinement, and validation statistics for *in situ* constructs.

**Table S3.** Simulation details for each simulated setup.

|  |  | Box Dim.<br>(nm) | No. Atoms | No. Solv | No. Ions |
| --- | --- | --- | --- | --- | --- |
| **Immature** | Monomer | 15.5 x 15.5 x 16.5 | 397,209 | 383,415 | 741 |
|  | Trimer-C3 | 15.5 x 15.5 x 16.5 | 386,172 | 375,309 | 747 |
|  | Hexamer-C6 | 15.0 x 15.0 x 16.0 | 345,563 | 322,799 | 647 |
|  | Lattice | 29.0 x 29.0 x 17.5 | 1,405,801 | 1,246,453 | 2,638 |
| **Hybrid** | Monomer | 14.3 x 14.7 x 15.2 | 300,210 | 296,416 | 577 |
|  | Trimer-C3 | 15.4 x 15.4 x 16.4 | 374,770 | 363,907 | 733 |
|  | Hexamer-C6 | 16.7 x 16.2 x 17.2 | 430,852 | 408,088 | 826 |
|  | Lattice | 29.3 x 29.3 x 20.5 | 1,677,546 | 1,518,198 | 3,492 |
| **Mature** | Monomer | 11.6 x 11.6 x 12.6 | 162,958 | 159,362 | 314 |
|  | Hexamer-C6 | 13.5 x 13.5 x 12.6 | 215,437 | 193,861 | 424 |

## References

1. Freed EO. HIV-1 assembly, release and maturation. Nature Reviews Microbiology. 2015/08/01 2015;13(8):484–496. doi:10.1038/nrmicro3490

2. Keller PW, Adamson CS, Heymann JB, Freed EO, Steven AC. HIV-1 maturation inhibitor bevirimat stabilizes the immature Gag lattice. J Virol. Feb 2011;85(4):1420–8. doi:10.1128/jvi.01926-10

3. Schur FK, Obr M, Hagen WJ, et al. An atomic model of HIV-1 capsid-SP1 reveals structures regulating assembly and maturation. Science. Jul 29 2016;353(6298):506–8. doi:10.1126/science.aaf9620

4. Wang D, Lu W, Li F. Pharmacological intervention of HIV-1 maturation. Acta Pharm Sin B. Nov 2015;5(6):493–9. doi:10.1016/j.apsb.2015.05.004

5. Wright ER, Schooler JB, Ding HJ, et al. Electron cryotomography of immature HIV-1 virions reveals the structure of the CA and SP1 Gag shells. The EMBO Journal. 2007;26(8):2218–2226. 10.1038/sj.emboj.7601664

6. Briggs JA, Riches JD, Glass B, Bartonova V, Zanetti G, Kräusslich HG. Structure and assembly of immature HIV. Proc Natl Acad Sci U S A. Jul 7 2009;106(27):11090–5. doi:10.1073/pnas.0903535106

7. Fuller SD, Wilk T, Gowen BE, Kräusslich HG, Vogt VM. Cryo-electron microscopy reveals ordered domains in the immature HIV-1 particle. Curr Biol. Oct 1 1997;7(10):729–38. doi:10.1016/s0960-9822(06)00331-9

8. Mendonça L, Sun D, Ning J, et al. CryoET structures of immature HIV Gag reveal six-helix bundle. Commun Biol. Apr 16 2021;4(1):481. doi:10.1038/s42003-021-01999-1

9. Fontana J, Keller PW, Urano E, Ablan SD, Steven AC, Freed EO. Identification of an HIV-1 Mutation in Spacer Peptide 1 That Stabilizes the Immature CA-SP1 Lattice. J Virol. Jan 15 2016;90(2):972–8. doi:10.1128/jvi.02204-15

10. Wagner JM, Zadrozny KK, Chrustowicz J, et al. Crystal structure of an HIV assembly and maturation switch. Elife. Jul 14 2016;5doi:10.7554/eLife.17063

11. Mallery DL, Kleinpeter AB, Renner N, et al. A stable immature lattice packages IP(6) for HIV capsid maturation. Sci Adv. Mar 2021;7(11)doi:10.1126/sciadv.abe4716

12. Renner N, Kleinpeter A, Mallery DL, et al. HIV-1 is dependent on its immature lattice to recruit IP6 for mature capsid assembly. Nat Struct Mol Biol. Mar 2023;30(3):370–382. doi:10.1038/s41594-022-00887-4

13. Mallery DL, Faysal KMR, Kleinpeter A, et al. Cellular IP(6) Levels Limit HIV Production while Viruses that Cannot Efficiently Package IP(6) Are Attenuated for Infection and Replication. Cell Rep. Dec 17 2019;29(12):3983–3996.e4. doi:10.1016/j.celrep.2019.11.050

14. Ricana CL, Lyddon TD, Dick RA, Johnson MC. Primate lentiviruses require Inositol hexakisphosphate (IP6) or inositol pentakisphosphate (IP5) for the production of viral particles. PLoS Pathog. Aug 2020;16(8):e1008646. doi:10.1371/journal.ppat.1008646

15. Dick RA, Zadrozny KK, Xu C, et al. Inositol phosphates are assembly co-factors for HIV-1. Nature. Aug 2018;560(7719):509–512. doi:10.1038/s41586-018-0396-4

16. Kleinpeter AB, Zhu Y, Mallery DL, et al. The Effect of Inositol Hexakisphosphate on HIV-1 Particle Production and Infectivity can be Modulated by Mutations that Affect the Stability of the Immature Gag Lattice. Journal of Molecular Biology. 2023/06/01/ 2023;435(11):168037. 10.1016/j.jmb.2023.168037

17. Ganser BK, Li S, Klishko VY, Finch JT, Sundquist WI. Assembly and Analysis of Conical Models for the HIV-1 Core. Science (New York, NY). 1999;283(5398):80–83. doi:doi:10.1126/science.283.5398.80

18. Highland CM, Tan A, Ricaña CL, Briggs JAG, Dick RA. Structural insights into HIV-1 polyanion-dependent capsid lattice formation revealed by single particle cryo-EM. Proceedings of the National Academy of Sciences. 2023;120(18):e2220545120. doi:doi:10.1073/pnas.2220545120

19. Briggs JA, Simon MN, Gross I, et al. The stoichiometry of Gag protein in HIV-1. Nat Struct Mol Biol. Jul 2004;11(7):672–5. doi:10.1038/nsmb785

20. Briggs JA, Grünewald K, Glass B, Förster F, Kräusslich HG, Fuller SD. The mechanism of HIV-1 core assembly: insights from three-dimensional reconstructions of authentic virions. Structure. Jan 2006;14(1):15–20. doi:10.1016/j.str.2005.09.010

21. Lanman J, Lam TT, Emmett MR, Marshall AG, Sakalian M, Prevelige PE, Jr. Key interactions in HIV-1 maturation identified by hydrogen-deuterium exchange. Nat Struct Mol Biol. Jul 2004;11(7):676–7. doi:10.1038/nsmb790

22. Meng X, Zhao G, Yufenyuy E, et al. Protease cleavage leads to formation of mature trimer interface in HIV-1 capsid. PLoS Pathog. 2012;8(8):e1002886. doi:10.1371/journal.ppat.1002886

23. Frank GA, Narayan K, Bess JW, Jr., et al. Maturation of the HIV-1 core by a non-diffusional phase transition. Nat Commun. Jan 8 2015;6:5854. doi:10.1038/ncomms6854

24. Woodward CL, Cheng SN, Jensen GJ. Electron cryotomography studies of maturing HIV-1 particles reveal the assembly pathway of the viral core. J Virol. Jan 15 2015;89(2):1267–77. doi:10.1128/jvi.02997-14

25. Ning J, Erdemci-Tandogan G, Yufenyuy EL, et al. In vitro protease cleavage and computer simulations reveal the HIV-1 capsid maturation pathway. Nat Commun. Dec 13 2016;7:13689. doi:10.1038/ncomms13689

26. Kucharska I, Ding P, Zadrozny KK, et al. Biochemical Reconstitution of HIV-1 Assembly and Maturation. J Virol. Feb 14 2020;94(5)doi:10.1128/jvi.01844-19

27. Wu C, Meuser ME, Rey JS, et al. Distinct Target Site of Lenacapavir in Immature HIV-1 and Concurrent Binding with the Maturation Inhibitor Bevirimat. Journal of the American Chemical Society. 2025/11/19 2025;147(46):42685–42700. doi:10.1021/jacs.5c13735

28. Cortines JR, Monroe EB, Kang S, Prevelige PE, Jr. A retroviral chimeric capsid protein reveals the role of the N-terminal β-hairpin in mature core assembly. J Mol Biol. Jul 22 2011;410(4):641–52. doi:10.1016/j.jmb.2011.03.052

29. Gross I, Hohenberg H, Huckhagel C, Kräusslich H-G. N-Terminal Extension of Human Immunodeficiency Virus Capsid Protein Converts the In Vitro Assembly Phenotype from Tubular to Spherical Particles. Journal of Virology. 1998;72(6):4798–4810. doi:10.1128/jvi.72.6.4798-4810.1998

30. Campbell S, Rein A. In vitro assembly properties of human immunodeficiency virus type 1 Gag protein lacking the p6 domain. J Virol. Mar 1999;73(3):2270–9. doi:10.1128/jvi.73.3.2270-2279.1999

31. Gross I, Hohenberg H, Wilk T, et al. A conformational switch controlling HIV - 1 morphogenesis. The EMBO Journal. 2000/01/04 2000;19(1):103–113-113. 10.1093/emboj/19.1.103

32. Campbell S, Fisher RJ, Towler EM, et al. Modulation of HIV-like particle assembly *in vitro* by inositol phosphates. Proceedings of the National Academy of Sciences. 2001;98(19):10875–10879. doi:doi:10.1073/pnas.191224698

33. Cook M, Freniere C, Wu C, Lozano F, Xiong Y. Structural insights into HIV-2 CA lattice formation and FG-pocket binding revealed by single particle cryo-EM. bioRxiv. Oct 9 2024;doi:10.1101/2024.10.09.617312

34. Arizaga F, Jr., Freniere C, Rey JS, et al. Exploring the Structural Divergence of HIV and SRLV Lentiviral Capsids. J Am Chem Soc. Sep 10 2025;147(36):32883–32895. doi:10.1021/jacs.5c09436

35. Kleinpeter AB, Freed EO. HIV-1 Maturation: Lessons Learned from Inhibitors. Viruses. Aug 26 2020;12(9)doi:10.3390/v12090940

36. Scholz I, Arvidson B, Huseby D, Barklis E. Virus particle core defects caused by mutations in the human immunodeficiency virus capsid N-terminal domain. J Virol. Feb 2005;79(3):1470–9. doi:10.1128/jvi.79.3.1470-1479.2005

37. Rossi E, Meuser ME, Cunanan CJ, Cocklin S. Structure, Function, and Interactions of the HIV-1 Capsid Protein. Life (Basel). Jan 29 2021;11(2)doi:10.3390/life11020100

38. Schirra RT, dos Santos NFB, Zadrozny KK, Kucharska I, Ganser-Pornillos BK, Pornillos O. A molecular switch modulates assembly and host factor binding of the HIV-1 capsid. Nature Structural & Molecular Biology. 2023/03/01 2023;30(3):383–390. doi:10.1038/s41594-022-00913-5

39. Stacey JCV, Tan A, Lu JM, James LC, Dick RA, Briggs JAG. Two structural switches in HIV-1 capsid regulate capsid curvature and host factor binding. Proceedings of the National Academy of Sciences. 2023;120(16):e2220557120. doi:doi:10.1073/pnas.2220557120

40. Jiang J, Ablan SD, Derebail S, et al. The interdomain linker region of HIV-1 capsid protein is a critical determinant of proper core assembly and stability. Virology. Dec 20 2011;421(2):253–65. doi:10.1016/j.virol.2011.09.012

41. Byeon IJ, Meng X, Jung J, et al. Structural convergence between Cryo-EM and NMR reveals intersubunit interactions critical for HIV-1 capsid function. Cell. Nov 13 2009;139(4):780–90. doi:10.1016/j.cell.2009.10.010

42. Wiegers K, Rutter G, Kottler H, Tessmer U, Hohenberg H, Kräusslich HG. Sequential steps in human immunodeficiency virus particle maturation revealed by alterations of individual Gag polyprotein cleavage sites. J Virol. Apr 1998;72(4):2846–54. doi:10.1128/jvi.72.4.2846-2854.1998

43. Mattei S, Tan A, Glass B, Müller B, Kräusslich HG, Briggs JAG. High-resolution structures of HIV-1 Gag cleavage mutants determine structural switch for virus maturation. Proc Natl Acad Sci U S A. Oct 2 2018;115(40):E9401–e9410. doi:10.1073/pnas.1811237115

44. Eisenstein F, Yanagisawa H, Kashihara H, Kikkawa M, Tsukita S, Danev R. Parallel cryo electron tomography on in situ lamellae. Nat Methods. Jan 2023;20(1):131–138. doi:10.1038/s41592-022-01690-1

45. Punjani A, Rubinstein JL, Fleet DJ, Brubaker MA. cryoSPARC: algorithms for rapid unsupervised cryo-EM structure determination. Nature Methods. 2017/03/01 2017;14(3):290–296. doi:10.1038/nmeth.4169

46. Zivanov J, Nakane T, Scheres SHW. Estimation of high-order aberrations and anisotropic magnification from cryo-EM data sets in RELION-3.1. IUCrJ. Mar 1 2020;7(Pt 2):253–267. doi:10.1107/s2052252520000081

47. Cheng J, Liu T, You X, et al. Determining protein structures in cellular lamella at pseudo-atomic resolution by GisSPA. Nat Commun. Mar 15 2023;14(1):1282. doi:10.1038/s41467-023-36175-y

48. Zheng W, Zhang Y, Wang J, et al. Visualizing the translation landscape in human cells at high resolution. Nat Commun. Nov 28 2025;16(1):10757. doi:10.1038/s41467-025-65795-9

49. Wang S, Brischigliaro M, Zhang Y, et al. Structural Basis of TACO1-Mediated Efficient Mitochondrial Translation. bioRxiv. 2025:2025.12.18.695269. doi:10.64898/2025.12.18.695269

50. Pettersen EF, Goddard TD, Huang CC, et al. UCSF Chimera--a visualization system for exploratory research and analysis. J Comput Chem. Oct 2004;25(13):1605–12. doi:10.1002/jcc.20084

51. Emsley P, Cowtan K. Coot: model-building tools for molecular graphics. Acta Crystallogr D Biol Crystallogr. Dec 2004;60(Pt 12 Pt 1):2126–32. doi:10.1107/s0907444904019158

52. Afonine PV, Poon BK, Read RJ, et al. Real-space refinement in PHENIX for cryo-EM and crystallography. Acta Crystallogr D Struct Biol. Jun 1 2018;74(Pt 6):531–544. doi:10.1107/s2059798318006551

53. Project CC. The CCP4 suite: programs for protein crystallography. *Acta crystallographica Section D*, Biological crystallography. 1994;50(Pt 5):760–763.

54. Agirre J, Atanasova M, Bagdonas H, et al. The CCP4 suite: integrative software for macromolecular crystallography. Acta Crystallogr D Struct Biol. Jun 1 2023;79(Pt 6):449–461. doi:10.1107/s2059798323003595

55. Burt A, Toader B, Warshamanage R, et al. An image processing pipeline for electron cryo-tomography in RELION-5. FEBS Open Bio. Nov 2024;14(11):1788–1804. doi:10.1002/2211-5463.13873

56. Zheng SQ, Palovcak E, Armache JP, Verba KA, Cheng Y, Agard DA. MotionCor2: anisotropic correction of beam-induced motion for improved cryo-electron microscopy. Nat Methods. Apr 2017;14(4):331–332. doi:10.1038/nmeth.4193

57. Rohou A, Grigorieff N. CTFFIND4: Fast and accurate defocus estimation from electron micrographs. J Struct Biol. Nov 2015;192(2):216–21. doi:10.1016/j.jsb.2015.08.008

58. Buchholz TO, Jordan M, Pigino G, Jug F. Cryo-CARE: Content-Aware Image Restoration for Cryo-Transmission Electron Microscopy Data. 2019:502–506.

59. Zheng S, Wolff G, Greenan G, et al. AreTomo: An integrated software package for automated marker-free, motion-corrected cryo-electron tomographic alignment and reconstruction. Journal of Structural Biology: X. 2022/01/01/ 2022;6:100068. 10.1016/j.yjsbx.2022.100068

60. Sofroniew N, Lambert T, Bokota G, et al. napari: a multi-dimensional image viewer for Python. Zenodo; 2024.

61. Gallay P, Hope T, Chin D, Trono D. HIV-1 infection of nondividing cells through the recognition of integrase by the importin/karyopherin pathway. Proc Natl Acad Sci U S A. Sep 2 1997;94(18):9825–30. doi:10.1073/pnas.94.18.9825

62. Aiken C, Trono D. Nef stimulates human immunodeficiency virus type 1 proviral DNA synthesis. J Virol. Aug 1995;69(8):5048–56. doi:10.1128/jvi.69.8.5048-5056.1995

63. Jennings J, Bracey H, Hong J, et al. The HIV-1 capsid serves as a nanoscale reaction vessel for reverse transcription. PLoS Pathog. Sep 2024;20(9):e1011810. doi:10.1371/journal.ppat.1011810

64. Webb B, Sali A. Comparative Protein Structure Modeling Using MODELLER. Curr Protoc Bioinformatics. Jun 20 2016;54:5.6.1–5.6.37. doi:10.1002/cpbi.3

65. Jorgensen WL, Chandrasekhar J, Madura JD, Impey RW, Klein ML. Comparison of simple potential functions for simulating liquid water. The Journal of Chemical Physics. 1983;79(2):926–935. doi:10.1063/1.445869

66. Humphrey W, Dalke A, Schulten K. VMD: visual molecular dynamics. Journal of molecular graphics. 1996;14(1):33–38.

67. Humphrey W, Dalke A, Schulten K. VMD: visual molecular dynamics. J Mol Graph. Feb 1996;14(1):33–8, 27-8. doi:10.1016/0263-7855(96)00018-5

68. Phillips JC, Hardy DJ, Maia JDC, et al. Scalable molecular dynamics on CPU and GPU architectures with NAMD. J Chem Phys. Jul 28 2020;153(4):044130. doi:10.1063/5.0014475

69. Huang J, Rauscher S, Nawrocki G, et al. CHARMM36m: an improved force field for folded and intrinsically disordered proteins. Nat Methods. Jan 2017;14(1):71–73. doi:10.1038/nmeth.4067

70. Best RB, Zhu X, Shim J, et al. Optimization of the additive CHARMM all-atom protein force field targeting improved sampling of the backbone φ, ψ and side-chain χ(1) and χ(2) dihedral angles. J Chem Theory Comput. Sep 11 2012;8(9):3257–3273. doi:10.1021/ct300400x

71. Mackerell AD, Jr., Feig M, Brooks CL, 3rd. Extending the treatment of backbone energetics in protein force fields: limitations of gas-phase quantum mechanics in reproducing protein conformational distributions in molecular dynamics simulations. J Comput Chem. Aug 2004;25(11):1400–15. doi:10.1002/jcc.20065

72. MacKerell AD, Bashford D, Bellott M, et al. All-atom empirical potential for molecular modeling and dynamics studies of proteins. J Phys Chem B. Apr 30 1998;102(18):3586–616. doi:10.1021/jp973084f

73. Elber R, Ruymgaart AP, Hess B. SHAKE parallelization. Eur Phys J Spec Top. Nov 1 2011;200(1):211–223. doi:10.1140/epjst/e2011-01525-9

74. Miyamoto S, Kollman PA. Settle: An analytical version of the SHAKE and RATTLE algorithm for rigid water models. Journal of Computational Chemistry. 1992;13(8):952–962. 10.1002/jcc.540130805

75. Essmann U, Perera L, Berkowitz ML, Darden T, Lee H, Pedersen LG. A smooth particle mesh Ewald method. The Journal of Chemical Physics. 1995;103(19):8577–8593. doi:10.1063/1.470117

76. Fiorin G, Klein ML, Hénin J. Using collective variables to drive molecular dynamics simulations. Molecular Physics. 2013/12/01 2013;111(22-23):3345–3362. doi:10.1080/00268976.2013.813594

77. Fiorin G, Marinelli F, Forrest LR, et al. Expanded Functionality and Portability for the Colvars Library. J Phys Chem B. Nov 14 2024;128(45):11108–11123. doi:10.1021/acs.jpcb.4c05604

78. Horn BKP. Closed-form solution of absolute orientation using unit quaternions. J Opt Soc Am A. 1987/04/01 1987;4(4):629–642. doi:10.1364/JOSAA.4.000629

79. Coutsias EA, Seok C, Dill KA. Using quaternions to calculate RMSD. J Comput Chem. Nov 30 2004;25(15):1849–57. doi:10.1002/jcc.20110

